# A Paracrine-to-Autocrine Shunt of GREM1 Fuels Colorectal Cancer Metastasis via ACVR1C

**DOI:** 10.1101/2025.07.11.664267

**Authors:** Huaixiang Zhou, Qunlong Jin, Zhang Fu, Yanming Yang, Yunfei Gao, Niu Wang, Bo Zhao, Long Gui, Jiang Li, Zijing Zhu, Ying Zhang, Yulong He, Ying Zhang, Shouqing Luo, Li Fu, Xudong Wu, Junjing Zhang, Xuetong Shen, Tao Wang, Youheng Jiang, Ningning Li

**Author notes:** Corresponding author: Ningning Li, Youheng Jiang, Tao Wang, Xuetong Shen. These authors contributed equally (Huaixiang Zhou, Qunlong Jin, Zhang Fu, Yanming Yang).

## Abstract

Tumor cells typically rely on paracrine stromal signals to guide malignant behaviors, but whether they acquire signaling autonomy to support metastasis remains unclear. We elucidate this in colorectal cancer (CRC) by uncovering a paracrine-to-autocrine shunt of Gremlin1 (GREM1), a canonical stromal-secreted antagonist of bone morphogenetic proteins (BMPs). We demonstrate that while GREM1 remains restricted to stromal cells in earlier-stage (I–III) CRC, its ectopic expression in tumor epithelium increases markedly in stage IV. Mechanistically, we identify ACVR1C as a novel, high-affinity epithelial receptor for GREM1. Their interaction activates SMAD2/3 signaling, which upregulates *SNAI1* and *GREM1*, establishing a feedback loop that amplifies epithelial-mesenchymal transition (EMT). Disrupting this loop impairs CRC metastasis *in vivo*. Clinically, epithelial GREM1 or ACVR1C expression predicts metastasis and poor survival. These findings define a paradigm in which tumor cells hijack stromal GREM1 to establish a GREM1–ACVR1C autocrine loop that sustains EMT and metastasis, marking a shift toward signaling autonomy and revealing a targetable vulnerability in advanced CRC.

## INTRODUCTION

Distant metastasis remains the leading cause of death in colorectal cancer (CRC) patients(Canellas-Socias et al., 2024). Epithelial–mesenchymal transition (EMT) is recognized as a critical early event initiating the metastatic cascade(Pastushenko and Blanpain, 2019). A complex interplay exists between tumor epithelial cells and various non-malignant components of the tumor microenvironment (TME), including cancer-associated fibroblasts (CAFs) and immune cells. This dynamic interaction is considered one of the central drivers of tumor initiation, progression, and phenotypic plasticity(de Visser and Joyce, 2023). Among these interactions, stroma-derived cytokines and chemokines help sustain the malignant phenotypes of tumor cells via paracrine signaling(Nee et al., 2023; Song et al., 2021; Tommelein et al., 2018). However, as tumor cells detach from the primary site to metastasize, they inevitably lose continuous support from the local TME—including stromal cells, extracellular matrix components, and localized signaling cues. Although tumor cells may, in certain contexts, co-migrate with CAFs and microenvironmental fragments into the circulation, the support they retain remains significantly limited compared to that at the primary site(Hurtado et al., 2020). Nonetheless, the support from these co-migrating components remains extremely limited compared to the primary site. This abrupt interruption of paracrine support is considered a major bottleneck that restricts the majority of tumor cells from successfully establishing distant colonies, with only a minority capable of seeding metastases eventually(Lambert et al., 2024).

As such, this phenomenon raises a fundamental yet unresolved question: how do a small subset of tumor cells maintain metastasis-associated phenotypes without sustained microenvironmental paracrine support? Given the extensive cross-talk between tumor cells and the TME, these metastasis-competent cells may have already undergone profound remodeling of their intrinsic properties via prolonged and dynamic interactions with TME at the primary site. This ability to sustain malignant behavior without external cues can be viewed as a form of ‘signaling autonomy,’ echoing the classic cancer hallmark of “self-sufficiency in growth signals” described by Hanahan and Weinberg(Hanahan and Weinberg, 2000). This hallmark refers to the ability of tumor cells to produce their own growth factors via autocrine loops, thereby bypassing the need for exogenous mitogenic stimulation and maintaining persistent signaling. While signaling autonomy has been extensively explored in the context of proliferation and survival, it is crucial to determine whether metastasizing tumor cells can achieve autonomous control over the programs that sustain EMT and migratory behavior—a capacity that could serve as an intrinsic driver of successful metastasis.

Within the TME, CAFs represent one of the most active stromal components and play essential roles in tumor initiation and progression through bidirectional interactions with cancer cells(Chen et al., 2021; Cords et al., 2024). Notably, within the highly heterogeneous CAF population(Luo et al., 2022), a GREM1-expressing subset has attracted increasing attention. Several studies have demonstrated that GREM1□ CAFs promote tumor progression via the paracrine secretion of GREM1 in both CRC(Jiang et al., 2024; Kobayashi et al., 2021; Li et al., 2025) and breast cancer(Ren et al., 2019a). Intriguingly, recent evidence indicates that GREM1 expression is not restricted to stromal CAFs. In pancreatic(Lan et al., 2022) and prostate cancers(Cheng et al., 2022), tumor epithelial cells have been shown to express and secrete GREM1, regulating their own phenotypic plasticity. Similarly, in Hereditary Mixed Polyposis Syndrome (HMPS), epithelial-specific upregulation of GREM1 is markedly enhanced in intestinal tissue(Jaeger et al., 2012a). Nonetheless, whether epithelial GREM1 expression occurs in sporadic CRC patients remains unclear, and its potential role in promoting cancer cell plasticity and metastasis has yet to be determined. Studies, including ours, have shown that modulating GREM1 expression in CRC cell lines can affect cell migration and EMT-related phenotypes(Li et al., 2022; Liu et al., 2019). Yet, whether CRC epithelial cells can “hijack” stromal GREM1 signals to ultimately activate their own GREM1 expression for malignant progression remains an open question. Mechanistically, GREM1 was initially identified as a canonical antagonist of the bone morphogenetic protein (BMP) signaling pathway(Brazil et al., 2015). Considering that BMPs are ligands rather than receptors and thus less likely to directly mediate GREM1’s potential autocrine function, it becomes critical to identify potential GREM1 receptors on CRC cells. Recent studies indicate that GREM1 also possesses cytokine-like functions, capable of binding to several noncanonical receptors, including VEGFR2(Mitola et al., 2010), FGFR1(Cheng et al., 2022), and EGFR(Park et al., 2020a), suggesting that it may influence cancer cell behavior through multiple signaling routes. However, it remains unclear whether GREM1 mediates its signaling autonomy during CRC progression via previously reported noncanonical receptors or other yet-to-be-identified ones.

This study aims to address two fundamental issues related to GREM1 signaling in CRC metastasis. First, we investigate whether CRC epithelial cells can acquire autonomous GREM1 signaling during tumor progression. Second, we examine whether GREM1 can regulate EMT and metastatic phenotypes through noncanonical receptors in CRC. By systematically elucidating how GREM1 signaling in CRC may undergo a paracrine-to-autocrine shunt, we seek to uncover a novel mechanism conferring signaling autonomy and metastatic competence. In doing so, our study positions GREM1 as an exemplar within a broader conceptual framework of metastasis wherein acquiring signaling autonomy is a critical driver of advanced progression. These efforts may not only provide a new paradigm for understanding how signaling autonomy contributes to metastasis but also establish a theoretical foundation for targeting the GREM1 signaling axis as a new anti-metastatic strategy.

## RESULTS

### Ectopic expression of GREM1 during CRC progression

In the gut, GREM1 marks a subpopulation of fibroblasts in both normal(Worthley et al., 2015) and tumor tissues(Kobayashi et al., 2021). Using *Grem1-CreER^T2^;Rosa-mTmG* mice(Worthley et al., 2015), we confirmed that Grem1^+^ cells are exclusively confined to the stromal compartment, and distributed along the intestinal isthmus and adjacent to α-SMA^+^ myofibroblasts(Kalluri and Zeisberg, 2006), two months post-tamoxifen (TMX) injection (Figures S1A–S1C). Extending this observation to humans, we found that GREM1 was likewise absent from epithelial cells in normal intestinal tissues, but sporadically expressed in stromal cells (Figure S1D), suggesting a conserved stromal specificity. Similarly, in human stage I-III CRC samples, GREM1 staining co-localized with VIMENTIN (VIM, a stromal cell marker, encoded by *VIM*)(Kalluri and Zeisberg, 2006; Pastushenko et al., 2018) and fibroblast activation protein (FAP, an activated fibroblast marker)(Kalluri and Zeisberg, 2006), but was mutually exclusive with β-CATENIN (β-CAT, a CRC cell marker, encoded by *CTNNB1*)(Zhao et al., 2022), CD68 (a macrophage marker)(Pastushenko et al., 2018), or α-SMA (a myofibroblast marker) (Figures S1E–S1J). These findings confirm that GREM1 is a bona fide stromal factor and GREM1^+^ stromal cells are a subtype of cancer-associated fibroblasts (CAFs), potentially contributing to CRC progression(Jiang et al., 2024; Kobayashi et al., 2021).

To systematically investigate the distribution and clinical significance of GREM1, we first performed immunohistochemical (IHC) staining on 106 human primary CRC samples spanning all four stages. We observed a stage-dependent redistribution of GREM1□ cells: in early-stage tumors (stage I–II), GREM1□ stromal cells were predominantly restricted to the peritumoral stroma; in stage III, these cells more frequently infiltrated the tumor parenchyma; in stage IV, infiltration of GREM1□ stromal cells showed a decreasing trend (with no statistically significant difference compared to stage III). Notably, in stage IV tumors, strong GREM1 staining emerged in a subset of tumor epithelial regions (Figures 1A–1C). These findings suggest a potential shift of GREM1 expression from stroma to epithelium during tumor progression.

**Figure 1.**
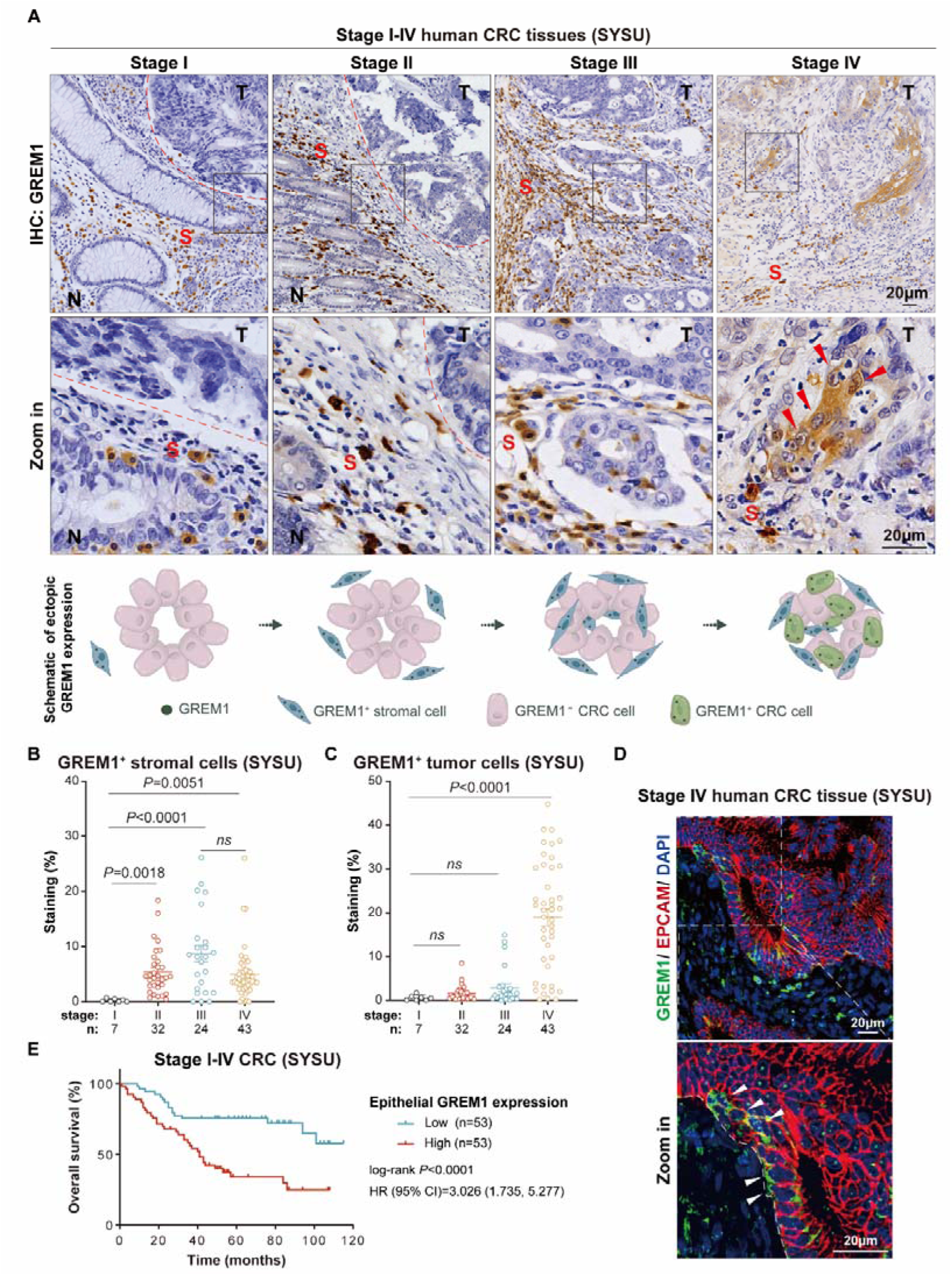
Ectopic expression of GREM1 protein in CRC. (**A**) Representative images of GREM1 immunohistochemical (IHC) staining (Upper) on human primary CRC samples and illustration of GREM1 expression pattern (Lower). The red dashed line separates tumor from adjacent normal area. N: normal areas; T: tumoral areas; S: stromal areas; Red arrow: GREM1^+^ CRC cells. (**B, C**) Quantification of GREM1□ stromal cell infiltration (B) and GREM1 expression in cancer cells (C) across stage I– IV primary CRC tissues from Sun Yat-sen University (SYSU). (**D**) Representative immunofluorescence (IF) images of GREM1 and EPCAM staining in stage IV human primary CRC tumors. A white dashed line separates epithelial and mesenchymal cells. White arrow: GREM1^+^ CRC cells. (**E**) Kaplan–Meier survival curves of 106 CRC patients from SYSU stratified by GREM1 levels in cancer cells. Patients were divided into high and low groups based on the cohort median. For B and C, data are mean ± s.e.m. *P* values were calculated using one-way ANOVA with Bonferroni multiple-comparison test (B, C). Significance was determined using a two-sided log-rank test. HR, hazard ratio (E). Scale bars, 20 μm.

To dissect the cellular sources and stage-specific dynamics of GREM1 expression at higher resolution, we analyzed publicly available single-cell RNA sequencing (scRNA-seq) datasets covering CRC samples from stages I–IV. Consistent with IHC data, GREM1 expression was primarily detected in fibroblasts and epithelial cells (Figures S2A–S2C). Stage-specific analysis revealed a marked upregulation of GREM1 in fibroblasts at stages III and IV (Figure S2D). Strikingly, GREM1□ epithelial cells were detected almost exclusively in stage IV tumors (Figure S2E), supporting the notion that ectopic GREM1 expression by tumor cells is a late event in CRC progression. This observation was further validated by colocalization of GREM1□ with an epithelial marker epithelial cell adhesion molecule (EPCAM)(Pastushenko et al., 2018) in tumor cells (Figure 1D). Finally, survival analysis demonstrated that patients with high GREM1 expression in tumor cells had significantly shorter overall survival compared to those with low expression (Figure 1E). Taken together, these results reveal that while GREM1□ CAFs are present throughout CRC progression, GREM1□ tumor epithelial cells emerge predominantly in advanced CRC, indicating that GREM1 expression, initially restricted to stromal cells, is progressively co-opted by tumor epithelial cells as CRC advances.

### ACVR1C is a novel GREM1 receptor in CRC

This spatiotemporal ectopic expression of GREM1 suggests that tumoral autocrine signaling may be initiated by preceding stromal paracrine cues. Given the potential for intercellular communication mediated by the infiltration of GREM1□ stromal cells into the tumor parenchyma, we hypothesized that CRC tumor cells might express GREM1 receptor(s), through which downstream signaling cascades drive tumoral GREM1 expression and promote CRC progression. To identify potential GREM1 receptors, we overexpressed HA-tagged GREM1 in the human CRC cell line HCT116 (Figure 2A). Mass spectrometry analysis of proteins pulled down using anti-HA beads identified activin A receptor type 1C (ACVR1C), a member of the TGFβ superfamily, as a potential GREM1 receptor (Figures 2B and S3A).

**Figure 2.**
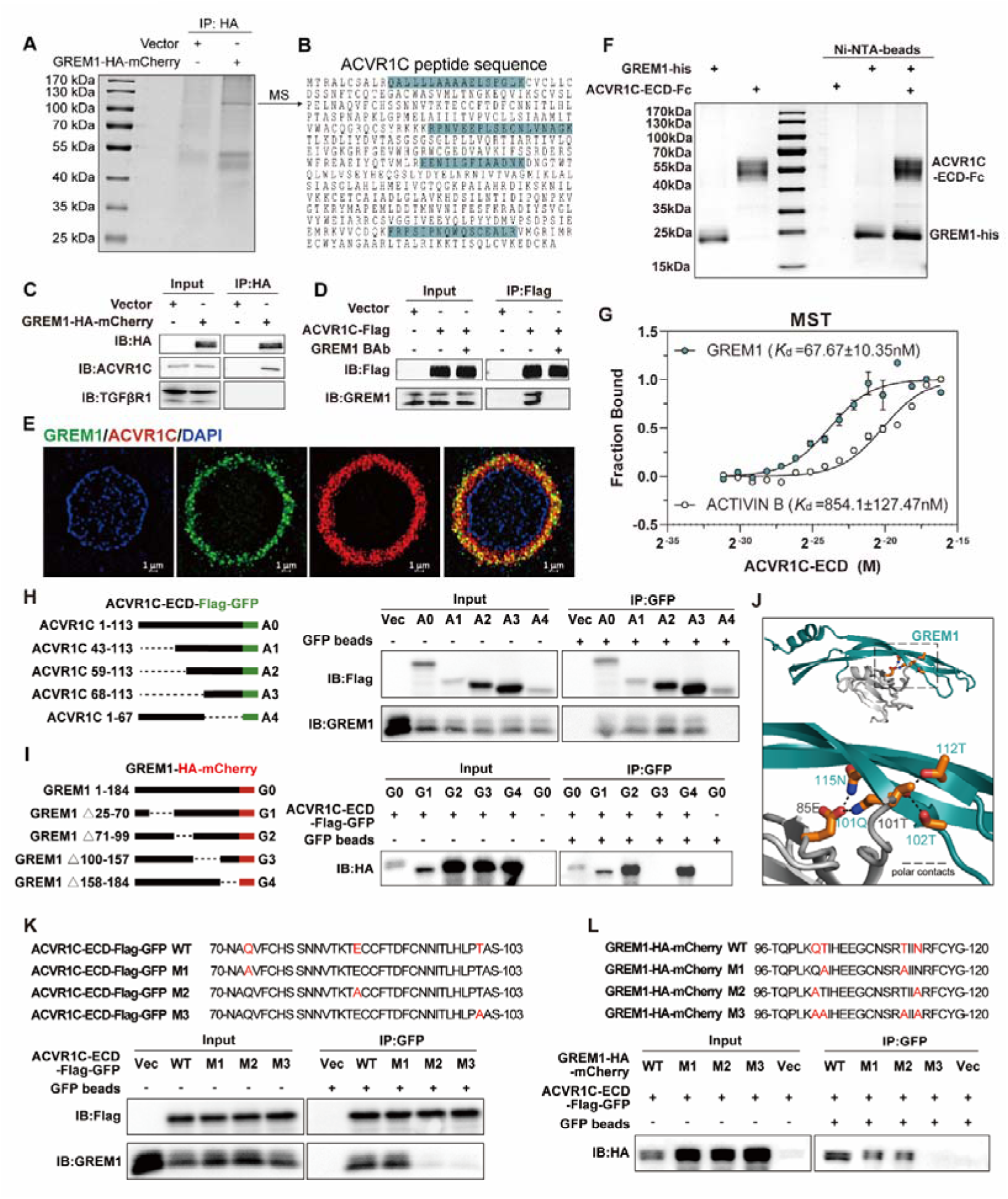
ACVR1C is a novel GREM1 receptor in CRC. (**A**) Proteins extracted from the HCT116 cells transfected with HA-mCherry-tagged-GREM1 were incubated with magnetic beads conjugated with an anti-HA antibody. Bound proteins were eluted and visualized by Coomassie Brilliant Blue staining. A protein band of ∼110 kDa was submitted for mass spectrometry (MS). (**B**) The full amino-acid sequence of human ACVR1C. The sequences in blue are the tryptic peptides identified by MS. (**C**) GREM1 coimmunoprecipitates with ACVR1C in HCT116 cells transfected with HA-mCherry-tagged GREM1 expressing plasmids. The bound proteins were immunoprecipitated with an anti-HA antibody and blotted by anti-HA, anti-ACVR1C, and anti-TGFβR1 antibodies. (**D**) ACVR1C coimmunoprecipitates with GREM1 in HCT116 cells transfected with Flag-tagged ACVR1C expressing plasmids. The bound proteins were immunoprecipitated with an anti-Flag antibody and blotted by an anti-GREM1 antibody. (**E**) Confocal microscopy images of GREM1 and ACVR1C in SW480 cells. Scale bars, 1 μm. (**F**) Interaction of purified GREM1 and ACVR1C-ECD protein demonstrated by pull-down experiments. (**G**) Increasing concentrations of recombinant ACVR1C-ECD-Fc protein (0-27.5 μM) were incubated with Red-labeled 50 nM recombinant GREM1-his or ACTIVIN B-his. MST was used to evaluate ACVR1C-ECD-Fc binding to GREM1-his or ACTIVIN B-his (n = 3 independent experiments), data are presented as mean ± s.e.m. (**H, I**) Diagrams of truncated ACVR1C (h, left) and truncated GREM1 (i, left), with corresponding coimmunoprecipitation results (right) comparing truncation mutants with binding partners. (**J**) Molecular docking of GREM1 and ACVR1C-ECD simulated by HDOCK (http://hdock.phys.hust.edu.cn/). Protein structures were generated from PDB (http://www.rcsb.org/, GREM1: 5AEJ) and AlphaFold (http://alphafold.ebi.ac.uk/, ACVR1C: Q8NER5). Docking module highlighting key amino acid residues in the binding pocket between GREM1 and ACVR1C. (**K**) Schematic of ACVR1C mutations (point mutations highlighted in red, Upper). Coimmunoprecipitation of ACVR1C and GREM1 is impaired by the AA_85_ mutant (M2) or the AA_101_ mutant (M3) of ACVR1C (Lower). (**L**) Schematic of GREM1 mutations (point mutations highlighted in red, Upper). Coimmunoprecipitation of ACVR1C and GREM1 is impaired by the AA_100-102/112/115_ mutant (M3) of GREM1 (Lower).

To validate this interaction, we performed co-immunoprecipitation (co-IP) assays using HA-tagged GREM1 in HCT116 cells. Immunoblotting revealed that GREM1 interacted with ACVR1C specifically, while no such interaction was detected with other members of the TGFβ superfamily such as TGFβR1 (Figure 2C). Similarly, co-IP assays using Flag-tagged ACVR1C confirmed the interaction with GREM1, which was abolished by a GREM1-blocking antibody (BAb) (Figure 2D). Confocal microscopy revealed the co-localization of ACVR1C with GREM1 in SW480 CRC cells (Figure 2E). We next examined whether a direct interaction exists between GREM1 and ACVR1C. We found that Fc-tagged ACVR1C extracellular domain (ACVR1C-ECD, AA_1-113_) and His-tagged full-length GREM1 were pulled down together, demonstrating a direct physical association between ACVR1C and GREM1 (Figure 2F). Further analysis of the binding affinity of GREM1 for ACVR1C using microscale thermophoresis (MST) revealed that ACVR1C-ECD exhibited a 12.6-fold higher affinity for GREM1 (*K*_d_ = 67.67 ± 10.35 nM) than that for ACTIVIN B (*K*_d_ = 854.1 ± 127.47 nM), a known ligand of ACVR1C(Tsuchida et al., 2004) (Figure 2G).

To further delineate the explicit interaction mode of GREM1 and ACVR1C, we constructed truncated GREM1 and ACVR1C-ECD. Co-IP assays showed that deletion of amino acids 100–157 (AA_100–157_) in GREM1 or 68–113 (AA_68–113_) in ACVR1C-ECD effectively abolished their interaction in HCT116 cells (Figures 2H and 2I). Based on these findings, we aimed to identify key residues mediating the interaction between GREM1 and ACVR1C. The HDOCK platform (http://hdock.phys.hust.edu.cn/) was utilized to simulate potential docking modalities between GREM1 (PDB: 5AEJ)(Kisonaite et al., 2016) and ACVR1C (AlphaFold prediction, https://alphafold.ebi.ac.uk/entry/Q8NER5) based on their structures. Residues Q101/T102/T112/N115 in GREM1 and Q72/E85/T101 in ACVR1C were predicted to be essential for binding (Figure 2J). To validate, we performed site-directed mutagenesis of the predicted residues to assess binding. Notably, the Q101A/T102A/T112A/N115A quadruple mutation in GREM1, or single mutations E85A or T101A in ACVR1C, significantly abrogated the GREM1-ACVR1C interaction (Figure 2K and 2L). Further, we found no detectable interaction between recombinant GREM1 and the ACVR1C-ECD double mutant (E85A/T101A) by assessing their binding affinity using MST (Figure S3B). These data suggest that Q101/T102/T112/N115 in GREM1, and E85/T101 in ACVR1C are key residues mediating their interaction. Clinically, IHC and scRNA-seq revealed that ACVR1C is expressed in tumor cells, with markedly elevated levels in stage IV CRC (Figures S3C–S3E). Notably, high ACVR1C expression correlates with poor prognosis in stage IV CRC patients, supporting ACVR1C’s tumor-promoting role and consistent with the clinical significance of GREM1 (Figure S3F). Taken together, these results demonstrate that ACVR1C is a novel receptor of GREM1 in CRC.

### Secretory GREM1 induces EMT via the ACVR1C-SMAD2/3 pathway but not TGF**β**R/BMPR pathways

To explore whether GREM1 serves as a functional ligand for ACVR1C in CRC cells, we first generated GREM1-enriched conditioned medium (GREM1-CM) and control conditioned medium (Vec-CM) using HEK293 cells (Figures S4A–S4C). We then performed RNA-sequencing (RNA-seq) on HCT116 cells treated with GREM1-CM or Vec-CM. Gene Set Enrichment Analysis (GSEA) revealed that SMAD2/3 and EMT pathways were significantly enriched in CRC cells treated with GREM1-CM (Figure 3A). ACVR1C is one of the receptors of the TGFβ superfamily, and it transduces signals primarily through the phosphorylation of SMAD2/3 (p-SMAD2/3)(Derynck and Zhang, 2003). Since commercial antibody for phosphorylated ACVR1C is not available, detection of p-SMAD2/3 serves as an effective proxy to reflect ACVR1C-SMAD2/3 activation. To confirm the effect of secretory GREM1 on the ACVR1C-SMAD2/3 pathway and EMT in CRC cells, we performed immunoblotting and found that GREM1-CM significantly increased p-SMAD2/3 levels in HCT116 and SW480 cells, which was effectively blocked by a GREM1 BAb (Figure 3B). Moreover, immunoblotting and RT–qPCR analyses also revealed that GREM1-CM induced significant downregulation of E-CADHERIN (E-CAD, encoded by *CDH1*)(Dongre and Weinberg, 2019) and upregulation of mesenchymal markers, including SNAIL (encoded by *SNAI1*), ZEB1, and β-CAT(Stemmler et al., 2019; Zhao et al., 2022) in SW480 and HCT116 cells. GREM1 BAb effectively blocked GREM1-CM-induced EMT activation (Figures 3B, S4D, and S4E). Considering that EMT serves as an effective mechanism through which tumor cells acquire stroma-like traits to promote invasion and metastasis, we tested whether blocking GREM1 could inhibit the invasive and migratory capacity of CRC cells. Indeed, *in vitro* scratch and transwell assays showed that GREM1-CM significantly enhanced migration and invasion abilities of HCT116 and SW480 cells. Remarkably, these effects were abolished by GREM1 BAb treatment (Figures S4F–S4I). These findings suggest that secretory GREM1 activates the ACVR1C-SMAD2/3 pathway and promotes EMT and subsequent cellular behavior, such as migration and invasion of CRC cells.

**Figure 3.**
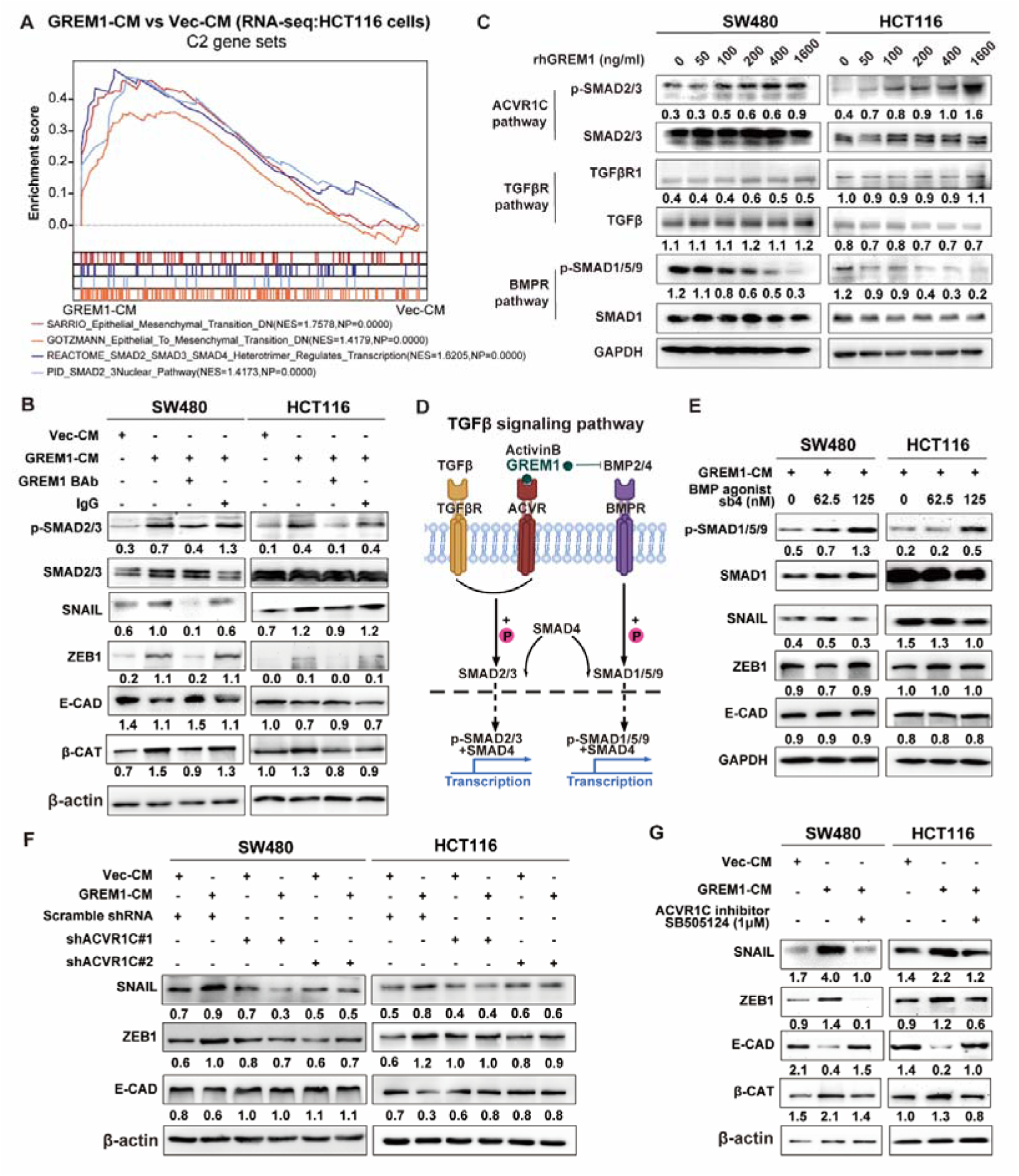
Secretory GREM1 binding of ACVR1C induced EMT via SMAD2/3 pathway. (**A**) RNA sequencing was performed on HCT116 cells treated with GREM1 conditioned medium (GREM1-CM) or control conditioned medium (Vec-CM), followed by GSEA of the C2 gene sets. NES, normalized enrichment score, all P-values equal to 0. (**B**) Activation levels of the ACVR1C pathway markers p-SMAD2/3 and SMAD2/3 and expression of epithelial marker E-CAD, mesenchymal markers SNAIL, ZEB1 and β-CAT were compared by immunoblotting analysis in SW480 and HCT116 cells treated with Vec-CM, GREM1-CM, GREM1-CM + GREM1 BAb or GREM1-CM + IgG. (**C**) Immunoblotting for the ACVR1C pathway markers p-SMAD2/3 and SMAD2/3 and the BMPR pathway markers p-SMAD1/5/9 and SMAD1 and the TGFβR pathway markers TGFβR1 and TGFβ from the lysates of HCT116 or SW480 cells with a concentration gradient of rhGREM1 treatment. (**D**) Schematic representation of the TGFβ signaling pathway, highlighting GREM1-induced activation of ACVR1C and inhibition of BMPR signaling. (**E**) Immunoblotting for the BMPR pathway markers p-SMAD1/5/9 and SMAD1 and expression of epithelial marker E-CAD, mesenchymal markers SNAIL and ZEB1 from the lysates of HCT116 or SW480 cells treated with GREM1-CM and different concentrations of BMP agonist, sb4. (**F**) SW480 and HCT116 cells were transfected with control shRNA (Scramble shRNA) or with one of two ACVR1C-targeting shRNAs (denoted shACVR1C#1 and shACVR1C#2) and separately treated with Vec-CM or GREM1-CM; epithelial marker E-CAD and mesenchymal markers SNAIL and ZEB1 were analyzed by immunoblotting. (**G**) Expression of epithelial markers E-CAD and mesenchymal markers SNAIL, ZEB1 and β-CAT were compared by immunoblotting analysis in SW480 and HCT116 cells treated with Vec-CM, GREM1-CM or GREM1-CM + SB505124 (1 μM). SB505124 inhibits the ACVR1C signaling pathway by impairing SMAD2/3 phosphorylation. CM, culture medium. For p-SMAD2/3, SMAD2/3 was used as a control. For other proteins, β-actin was used as a loading control.

The TGFβ superfamily signals through BMPRs, ACVRs, and TGFβRs. Although both ACVRs and TGFβRs converge on the SMAD2/3 axis(Derynck and Zhang, 2003), and GREM1-CM robustly activated TGFβ superfamily signaling in HEK293 cells (Figure S4J), our co-IP analysis showed no direct interaction between GREM1 and

TGFβR1 (Figure 2C). To determine whether GREM1 activates SMAD2/3 via TGFβR1 or ACVR1C, we treated SW480 and HCT116 cells with increasing concentrations of recombinant human GREM1 (rhGREM1). This led to a dose-dependent increase in p-SMAD2/3 (Figure 3C), suggesting activation of a SMAD2/3-coupled receptor. To exclude the possibility that GREM1 indirectly stimulates TGFβ signaling, we examined whether rhGREM1 alters the expression of TGFβ or TGFβR1 in CRC cells. No changes were observed across all doses tested (Figure 3C), indicating that GREM1 does not upregulate endogenous TGFβ signaling components. Together, these data suggest that GREM1-induced SMAD2/3 activation is unlikely to be mediated by TGFβR1 and instead proceeds via the ACVR1C pathway (Figure 3D). Considering that GREM1 is a canonical antagonist of BMP and that BMP receptors (BMPRs) exert their function through the phosphorylation of SMAD1/5/9 (p-SMAD1/5/9), we sought to investigate whether GREM1 regulates EMT via BMPR superfamily pathways. As expected, rhGREM1 suppressed the p-SMAD1/5/9 (Figure 3C), in keeping with the canonical role of GREM1 as a BMP inhibitor. However, rescue of BMP signaling using the specific agonist sb4, which exclusively increases p-SMAD1/5/9 levels without affecting p-SMAD2/3(Bradford et al., 2019), failed to reverse GREM1-induced EMT marker changes (Figure 3E). This definitive exclusion of BMPR-SMAD1/5/9 involvement establishes that GREM1 promotes EMT independently of its classical role as a BMP inhibitor.

Subsequently, to examine whether GREM1 promotes EMT through activation of the ACVR1C-SMAD2/3 pathway, we either stably knocked down ACVR1C (shACVR1C) or inhibited SMAD2/3 phosphorylation using SB505124(DaCosta Byfield et al., 2004). Our immunoblotting and RT–qPCR analysis revealed that shACVR1C or SB505124 significantly blocked GREM1-CM-induced changes in EMT marker expression (*i.e.* E-CAD, ZEB1, β-CAT and SNAIL), indicating that ACVR1C-SMAD2/3 activation is required for GREM1-driven EMT in SW480 and HCT116 CRC cells (Figures 3F, 3G, S5A, S5B, S6A, and S6B). In addition, GREM1-CM-induced invasion and migration of these CRC cells was abolished by shACVR1C or SB505124 (Figures S5C–S5G and S6C–S6F). Collectively, these findings demonstrate that GREM1 induces EMT, as well as subsequent migration and invasion, by activating the ACVR1C-SMAD2/3 pathway.

p-SMAD2/3 form a complex with SMAD4 that translocates into the nucleus and acts as a transcriptional regulator(Shi and Massagué, 2003). *SNAI1* has been identified as a transcriptional target of the SMAD2/3/4 complex in several cancers, including CRC(Asnaghi et al., 2019; Cai et al., 2019; Yang et al., 2021). To define the direct binding sites involved in this regulation in CRC cells, we queried the JASPAR database (http://jaspar.genereg.net/) and identified three candidate SMAD2/3/4 binding sites (–967, –787, and –186) within the *SNAI1* promoter (Figure S6G). Chromatin immunoprecipitation (ChIP) followed by qPCR confirmed specific binding of SMAD2/3/4 to the –787 site. Importantly, inhibition of ACVR1C with SB505124 significantly decreased this binding (Figure S6H). Overall, our data reveal that secretory GREM1 is a specific functional ligand that activates the ACVR1C– SMAD2/3–SNAIL signaling axis, thereby promoting EMT, invasion and migration of CRC cells.

### Exogenous GREM1 induction of endogenous *GREM1* transcription reinforces EMT in CRC cells via the ACVR1C-SMAD2/3 pathway

Our data above show that there is a marked increase in the proportion of GREM1^+^ epithelial tumor cells in stage IV CRC (Figures 1C and S2E). To determine whether exogenous GREM1 can regulate endogenous GREM1 expression, we treated SW480 cells with increasing concentrations of rhGREM1. RT-qPCR and immunoblotting analyses revealed a dose-dependent upregulation of tumor GREM1 expression in response to rhGREM1 stimulation (Figures 4A–4C). However, the mechanism by which exogenous GREM1 triggers endogenous *GREM1* transcription in advanced CRC cells remains unclear. Notably, we also observed that ACVR1C expression was markedly upregulated in stage IV CRC (Figures S3D and S3E), coinciding with the emergence of epithelial GREM1 expression, whereas stromal GREM1 was already abundantly present in both stage III and IV tumors (Figures 1B, 1C, S2D, and S2E). This spatiotemporal concordance suggests that epithelial GREM1 induction may depend on elevated ACVR1C expression. To investigate the role of the ACVR1C-SMAD2/3 pathway in the exogenous GREM1-mediated tumor *GREM1* transcription, we either overexpressed ACVR1C or inhibited the pathway using SB505124 in SW480 cells following rhGREM1 treatment. Interestingly, ACVR1C overexpression further enhanced endogenous *GREM1* expression, whereas SB505124 treatment completely reversed the effect of rhGREM1 (Figure 4D). To investigate whether SMAD2/3/4 act as transcription factors for *GREM1*, we utilized the JASPAR database and identified five candidate SMAD2/3/4 binding sites (-733, -612, -446, -316, and -3) in the *GREM1* promoter region (Figure 4E). To validate the predicted results, we performed a ChIP–qPCR analysis, which revealed that SMAD2/3/4 bound the *GREM1* promoter at the -733 and -612 sites. Notably, SB505124 treatment significantly reduced this binding (Figure 4F). These data demonstrate that exogenous GREM1 efficiently induces endogenous *GREM1* transcription in CRC cells via the ACVR1C-SMAD2/3 signaling pathway.

**Figure 4.**
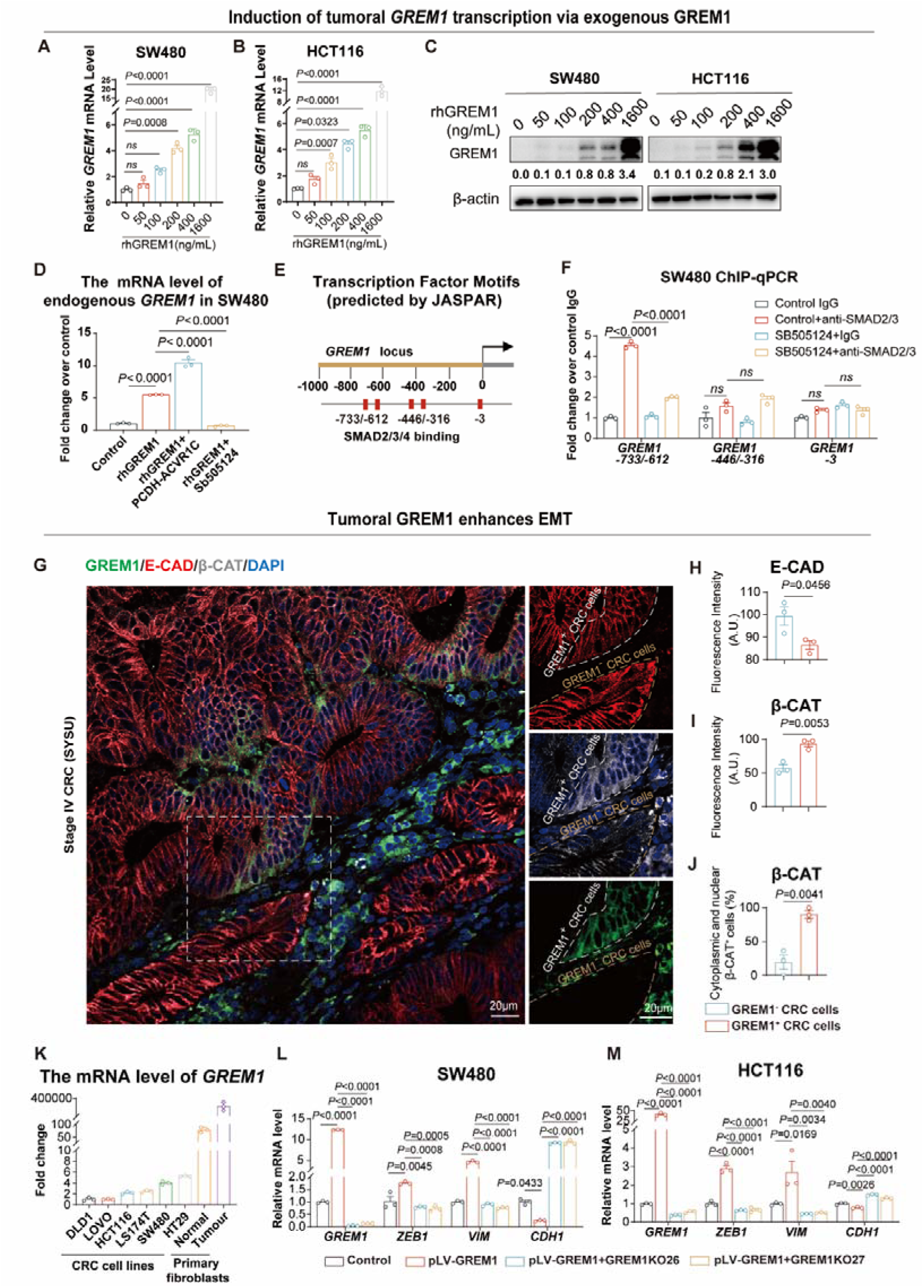
Secreted GREM1 upregulates *GREM1* transcription via ACVR1C-SMAD2/3 pathway to reinforce EMT in CRC. **(A, B**) RT–qPCR analysis of *GREM1* mRNA levels in SW480 (A) and HCT116 (B) cells treated with increasing concentrations of rhGREM1. n = 3 biological replicates. (**C**) Immunoblotting of GREM1 protein levels in SW480 and HCT116 cells treated with the concentration gradient of rhGREM1 used in (A) and (B). (**D**) RT–qPCR analysis of *GREM1* expression in SW480 cells treated with rhGREM1 (400 ng/mL). n = 3 biological replicates. (**E**) SMAD2/3/4 binding motifs in the *GREM1* promoter, predicted by the JASPAR database. Red boxes denote predicted binding sites upstream of the transcription start site (TSS). (**F**) ChIP analysis of p-SMAD2/3 binding sites on the *GREM1* promoter in SW480 cells, treated as indicated. IgG control or SMAD2/3 antibodies were used for ChIP and DNA was quantified by qPCR. DNA levels for each condition were normalized to the input, and the fold-change was calculated over the vehicle control (n = 3 independent experiments). *ns*: not significant. (**G**) Representative IF images for staining of GREM1, E-CAD and β-CAT in stage □ human primary CRC tumors from SYSU. The white square-dashed box in the left image indicates the location of the corresponding region in the right image. The white irregular-dashed box marks the GREM1□ CRC cells, while the brown irregular-dashed box marks the GREM1^−^ CRC cells. Scale bars, 20 μm. (**H, I)** Quantification of the fluorescence intensity of E-CAD (H) and β-CAT (I) between GREM1^+^ and GREM1^−^ CRC subpopulations shown in (G) (n = 3 patients). (**J**) Quantification of the percentage of cells exhibiting cytoplasmic and nuclear translocation of β-CAT in the conditions shown in (G) (n = 3 patients). (**K**) RT–qPCR analysis of *GREM1* mRNA expression levels in CRC cell lines (DLD1, LOVO, HCT116, LS174T, SW480, and HT29), primary fibroblasts (normal fibroblasts and tumoral fibroblasts) derived from the gut of human CRC patients. (**L, M**) RT–qPCR analysis of *GREM1*, epithelial marker *CDH1* and mesenchymal markers *ZEB1* and *VIM* in HCT116 cells infected with control (pLV) or GREM1-overexpressing (pLV-GREM1) lentiviruses or GREM1-overexpressing cells transduced with one of two GREM1-targeting sgRNAs (denoted GREM1KO26 and GREM1KO27), n = 3 independent experiments. For A, B, D, F and H–M, data are presented as mean ± s.e.m. *P* values were calculated using one-way ANOVA with Bonferroni multiple-comparison test (A, B, D, F, L, M) and two-tailed Student’s t-test (H–J).

Having established that exogenous GREM1 promotes EMT in CRC cells, we sought to determine whether endogenous GREM1 exerts a similar function. In normal epithelial cells, β-CAT is localized at the cellular membrane with the adhesion molecule E-CAD(Clevers, 2006). However, during tumor progression and the onset of EMT, E-CAD is gradually lost, and a portion of β-CAT translocates to the cytoplasm and nucleus(Inomata et al., 1996; Sánchez-Tilló et al., 2011). To delineate the bona fide correlation between epithelial GREM1 expression and EMT hallmarks in CRC, we performed immunofluorescence (IF) staining on stage IV CRC clinical samples, which contained both GREM1^+^ and GREM1^−^ CRC cells. Remarkably, GREM1^+^ CRC cells exhibited a significant loss of E-CAD, along with increased β-CAT expression and its translocation to the cytoplasm and nucleus, compared to adjacent GREM1^−^counterparts within the same tumor (Figures 4G–4J).

We previously found that forced *GREM1* expression (pLV-GREM1) in CRC cells enhanced their EMT and metastatic traits(Li et al., 2022). Given the typically low expression of GREM1 in CRC cell lines compared to normal and tumoral fibroblasts (Figure 4K), we sought to model the therapeutic blockade of endogenous GREM1 in advanced tumors. To this end, we used CRISPR/Cas9 techniques to knock out *GREM1* in SW480 and HCT116 cells that stably expressed pLV-GREM1. RT–qPCR analysis revealed that pLV-GREM1 resulted in a significant change in EMT markers, including a marked decrease in *CDH1* and an increase in *SNAI1, VIM,* and *ZEB1,* while *GREM1* knockout significantly restored the expression of these markers, indicating that endogenous GREM1 promotes EMT within CRC cells (Figures 4L and 4M). Collectively, these data demonstrate that exogenous GREM1 induces endogenous *GREM1* transcription in CRC cells through the ACVR1C–SMAD2/3 pathway, establishing a self-propelling loop that promotes EMT. Given our observation of a reduction in GREM1□ stromal cells in stage IV CRC compared to stage III (Figure 1B), we speculate that this self-sustaining GREM1 feedback loop enables CRC cells to maintain EMT independently of stromal inputs, reducing their reliance on stromal GREM1. This autonomous signaling cascade comprises multiple nodes that may be amenable to therapeutic intervention aimed at halting CRC metastasis.

### Targeting the GREM1-ACVR1C axis to inhibit EMT and metastasis of CRC *in vivo*

To validate the GREM1-ACVR1C-induced autocrine GREM1 feedback loop, we applied genetic and pharmacological strategies (Figure S6I). First, we evaluated the impact of stromal paracrine GREM1 on CRC *in vivo*. To block exogenous GREM1, we crossed *Grem1-CreER^T2^;Rosa-LSL-DTA* mice with *APC^Min/+^* mice(Moser et al., 1990), a well-established model for CRC proliferation and EMT that develops intestinal tumors by 10 weeks(Babaei-Jadidi et al., 2011; Ren et al., 2019b; Sánchez-Tilló et al., 2011), generating AGD mice for TMX-induced depletion of GREM1^+^ stromal cells (Figure 5A). Intriguingly, ablation of Grem1*^+^* stromal cells for 6 weeks postnatally did not significantly alter the number or size of intestinal tumors in *APC^Min/+^* mice (Figure S7A and S7B). Subsequently, we delved deeper into whether loss of paracrine GREM1 could restrain the malignant potential of intestinal tumor cells. As expected, IF staining in the GREM1*^+^* stromal cell infiltration zone of the *APC^Min/+^* intestines revealed a marked loss of E-cad in tumor cells (Figure 5B, upper left panels), accompanied by increased β-cat expression and its translocation to the cytoplasm and nucleus (Figure 5B, upper right panels). In contrast, upon depletion of GREM1*^+^* stromal cells in AGD mice, E-cad expression was significantly elevated (Figure 5C), with intense and continuous localization along the tumor cell membrane (Figure 5B, bottom left). In parallel, β-cat staining was reduced and restricted to the cell membrane (Figures 5D and 5E), co-localizing with E-cad, and rarely observed in the cytoplasm or nucleus (Figure 5B, bottom right panels). These results suggest a critical role for exogenous Grem1 in orchestrating malignant cell behaviors. Next, to evaluate the impact of paracrine Grem1 on CRC metastasis, we injected luciferase-labeled murine rectal cancer cells (MC38-luc) into Grem1*^+^* cell-depleted (GD) or control mice (Figures S7C and S7D). Cells were administered via the tail vein to induce lung metastasis, or into the spleen or cecum wall to induce liver metastasis(Zhang et al., 2021) (Figures S7E–S7G). Strikingly, we observed a significant reduction in lung and liver metastases of CRC cells in GD mice compared with *Grem1-CreER^T2^*or *Rosa-LSL-DTA* controls (Figures 5F, 5G, and S7H–S7K), suggesting that the stromal factor Grem1 is vital for CRC metastasis.

**Figure 5.**
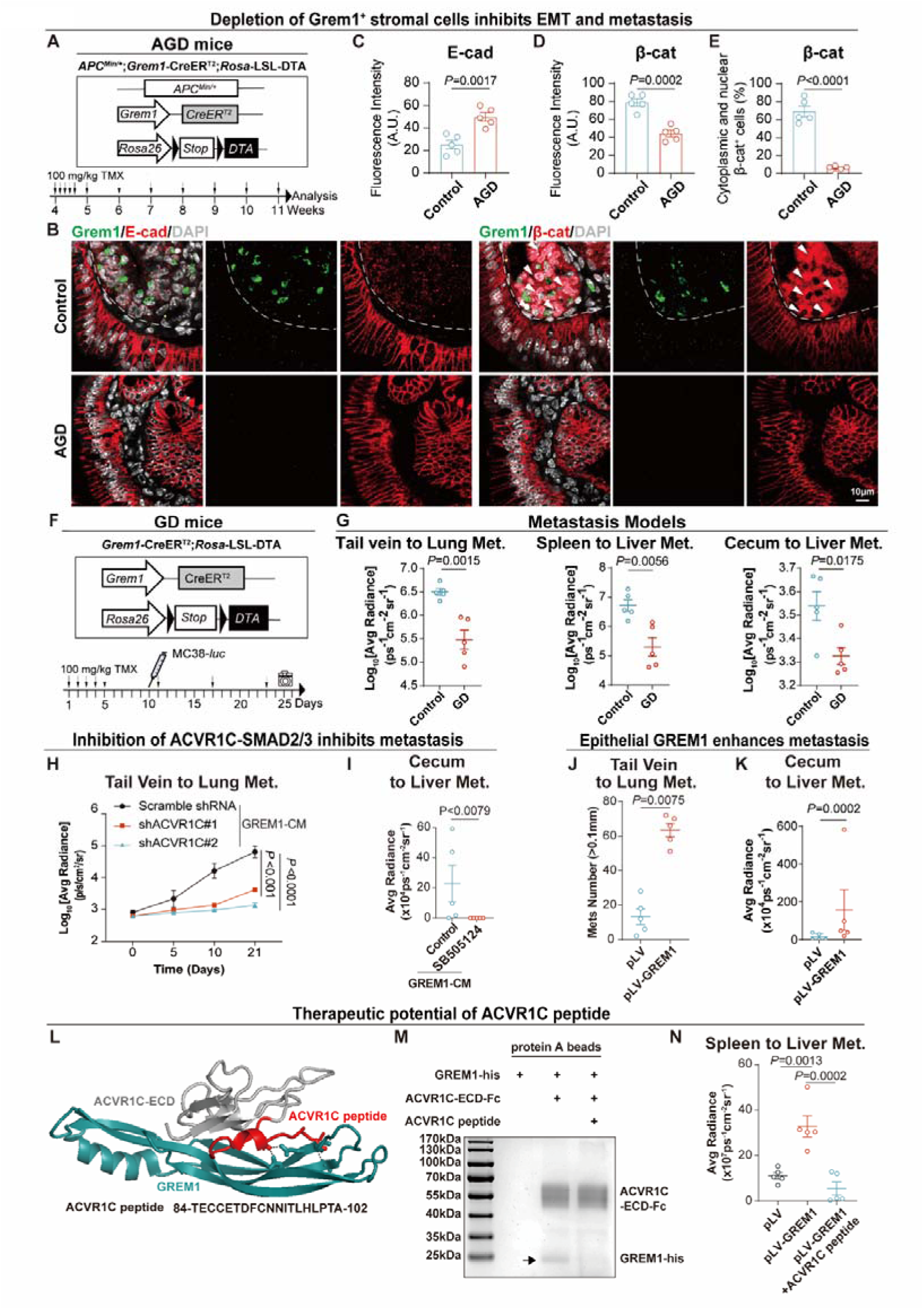
Targeting the GREM1-ACVR1C axis to inhibit EMT and metastasis of CRC *in vivo*. (**A**) Diagram of the AGD Mouse Model (*APC^Min/+^;Grem1-CreER^T2^;Rosa-LSL-DTA*) and the experimental approach. Grem1□ cells are selectively deleted through tamoxifen (TMX)-induced DTA expression via Cre-lox recombination. Mice received TMX (100 mg/kg) by oral gavage for five consecutive days starting at 4 weeks of age, followed by weekly TMX administration until analysis at 11 weeks. The black triangle indicates the loxP sites. (**B**) Representative IF images of staining for Grem1 and E-cad and β-cat in AGD or control mice (*APC^Min/+^;Grem1-CreER^T2^*or *APC^Min/+^;Rosa-LSL-DTA*). The white irregular dashed box indicates the infiltrated region of Grem1^+^ stromal cells. Scale bars, 10 μm. (**C, D**) Quantification of the fluorescence intensity of E-cad (C) and β-cat (D) between AGD and control mice shown in (B) (n = 5 mice per group). (**E**) Quantification of the percentage of cells exhibiting cytoplasmic and nuclear translocation of β-cat under the conditions shown in (B) (n = 5 mice per group). (**F**) Diagram of the GD mouse CRC model and the experimental approach. GD and control (*Grem1-CreER^T2^*or *Rosa-LSL-DTA*) mice received TMX (100 mg/kg) by oral gavage for five consecutive days starting at 6 weeks of age, followed by weekly doses. At designated time points, MC38-luciferase (MC38-luc) cells were injected intravenously, intrasplenically, or into the cecal wall. Lung and liver metastases were imaged and quantified every five days using the IVIS Lumina Imaging System. The black triangle indicates the loxP sites. (**G**) Quantification of lung and liver metastases. GD or control mice were injected intravenously, intrasplenically, or into the cecum wall with MC38-luc cells. Lung or liver metastases were imaged and quantified by IVIS Lumina Imaging System and quantified (n = 5 mice per group). met., metastases. (**H**) Quantification of the lung metastases. Mice were injected intravenously with HCT116□luciferase (HCT116□luc) cells, which were transduced with Scramble shRNA or shACVR1C and treated with GREM1-CM. Lung metastases were imaged and quantified by IVIS Lumina Imaging System. (**I**) Quantification of liver metastases. The cecum of NOG mice was injected with HCT116□luc cells with GREM1-CM treatment, and mice were treated with SB505124 or a solvent control via abdominal injection. Liver metastases were imaged and quantified by IVIS Lumina Imaging System (n = 5 mice per group). (**J**) Quantification of lung metastases. Mice were injected intravenously with HCT116 cells infected with pLV or pLV-GREM1 lentivirus. Lung tissue sections were stained for Ki67 by IHC, and lung metastases larger than 0.1 mm were counted (n = 5 mice per group). (**K**) Quantification of liver metastases. NOG mice’s cecum wall was injected with HCT116□luc cells (pLV/ pLV-GREM1). Liver metastases were imaged and quantified by IVIS Lumina Imaging System (n = 5 mice per group). (**L**) Docking module of GREM1 and ACVR1C-ECD, highlighting key amino acid sequence between GREM1 and ACVR1C (ACVR1C peptide). (**M**) The pull-down assay confirmed that ACVR1C peptide blocked the interaction between GREM1 and ACVR1C-ECD protein. The black arrow indicates the GREM1-his band. (**N**) Quantification of liver metastases. NOG mice’s spleen was injected with HCT116□luc cells (pLV/ pLV-GREM1). Treatment was started 48 h after cell injection. ACVR1C peptide was given once every other day (10□mg/kg i.v.). i.v.: intravenous injection. Liver metastases were imaged and quantified by IVIS Lumina Imaging System (n = 5 mice per group). For C–E, G–K, and N, data are presented as mean ± s.e.m. *P* values were calculated using two-tailed Student’s t-test (C–E, G, I–K) or one-way ANOVA with Bonferroni multiple-comparison test (H, N).

To determine the role of ACVR1C and its downstream SMAD2/3 pathway in GREM1-mediated EMT *in vivo*, we first pre-treated HCT116 cells carrying a luciferase reporter (HCT116-luc), stably expressing either shACVR1C or scramble shRNA, with GREM1-CM, followed by subcutaneous transplantation into nude mice. Notably, ACVR1C knockdown significantly inhibited subcutaneous tumor growth (Figures S8A–S8C), accompanied by a significant increase in epithelial gene expression (e.g. *CDH1*) and a marked decrease in mesenchymal gene expression (e.g. *SNAI1, CTNNB1,* and *ZEB1*), as shown by RT–qPCR analysis (Figure S8D). These findings were further corroborated by IF staining of EMT markers, including E-CAD and SNAIL (Figures S8E–S8G). Meanwhile, we also harnessed SB505124 as a pharmacological alternative to knock down ACVR1C. Consistent with the genetic approach, SB505124 treatment yielded similar effects on local tumor progression, further substantiating the contextual role of the ACVR1C-SMAD2/3 pathway (Figures S9A–S9G). Importantly, to investigate the impact of the ACVR1C-SMAD2/3 axis on metastasis, we injected GREM1-CM-pre-treated HCT116-luc cells, stably expressing either shACVR1C or scramble shRNA, into the tail vein of nude mice. Remarkably, lung metastasis was profoundly suppressed in the *ACVR1C* knockdown group compared with controls (Figures 5H and S8H). In parallel, we inoculated GREM1-CM-pre-treated HCT116-luc cells into the cecum wall of NOG (NOD/Shi-scid/IL-2Rγ) mice, a new generation of severely immunodeficient mice(Ito et al., 2002). We found that SB505124 treatment resulted in a significant reduction in liver metastasis (Figures 5I and. S9H). Collectively, these findings establish the ACVR1C-SMAD2/3 axis as a critical effector of stroma-derived GREM1, and reveal that its inhibition provides an effective strategy to counteract GREM1-induced EMT and metastasis in CRC *in vivo*.

Further, to examine the contribution of tumor-autocrine GREM1 to EMT and metastasis of CRC *in vivo*, we first inoculated HCT116 cells carrying pLV-GREM1 into nude mice, which led to significantly enhanced subcutaneous tumor growth (Figures S10A–S10C). Subsequent RT–qPCR analysis demonstrated that autocrine GREM1 upregulated *SNAI1, VIM,* and *ZEB1* while downregulating *CDH1* (Figure S10D). These findings were confirmed by IF staining for EMT markers, including E-CAD and SNAIL (Figures S10E–S10G), supporting that autocrine GREM1 enhances the EMT process in CRC cells. Next, to confirm the role of tumoral GREM1 in enhancing CRC metastasis, we conducted metastasis assays by injecting HCT116-luc cells expressing either pLV-GREM1 or the control vector via the tail vein or into the cecum wall. We found that overexpression of tumoral GREM1 resulted in a significant increase in lung and liver metastasis compared with controls (Figures 5J, 5K, S10H, and S10I). In summary, these findings demonstrate that tumoral GREM1 promotes both EMT and metastasis of CRC cells *in vivo*.

### Targeting the GREM1-ACVR1C interaction interface to inhibit metastasis of CRC *in vivo*

In clinical research, the molecular complexity and shared signaling of CRC limit the effective targeted therapy. While the TGFβ superfamily is a promising therapeutic axis(Jung et al., 2017), its intertwined and overarching roles in development and physiology make selective targeting difficult without systemic toxicity(Liu et al., 2021; Peng et al., 2022). Targeting protein–protein interaction (PPI) interfaces is thought to further enhance specificity and reduce off-target effects(Modell et al., 2016). Our investigation of the GREM1-ACVR1C interaction unveiled a potential avenue to overcome these limitations. Based on the principle of interface disruption, we designed a peptide inhibitor derived from amino acid residues 84-102 (AA84-102) of ACVR1C (hereafter referred to as the ACVR1C peptide). We further examined the binding affinity between GREM1 and the ACVR1C peptide using MST. We found that GREM1 exhibited an affinity for the ACVR1C peptide comparable to that for the ACVR1C-ECD (*K*_d_ = 92.30 ± 7.51 nM) (Figure S10J). Pull-down assays showed that the introduction of ACVR1C peptide significantly blocked the GREM1-ACVR1C binding (Figures 5L and 5M), suggesting that the peptide disrupts the GREM1-ACVR1C interaction by potently and competitively binding to GREM1. Further, we employed a spleen-to-liver metastasis model to evaluate the functional effect of the ACVR1C peptide in CRC metastasis. Notably, administration of the ACVR1C peptide effectively suppressed the increase in liver metastasis induced by tumor-specific GREM1 overexpression (Figures 5N and S10K), suggesting that our ACVR1C peptide significantly attenuated CRC progression by blocking the metastasis-promoting effects of tumor-autonomous autocrine GREM1-ACVR1C signaling, laying the foundation for the development of a novel class of peptide-based targeted therapies in CRC.

## DISCUSSION

Whether tumor cells can internalize stromal paracrine signals and convert them into autocrine loops during metastasis has remained unclear. Here, we show that in advanced colorectal cancer, GREM1 undergoes a stromal-to-epithelial shunt and binds ACVR1C to activate SMAD2/3, inducing both *SNAI1* and *GREM1* expression. This establishes a self-amplifying autocrine circuit that drives EMT, conferring signaling autonomy and promoting metastasis. A functional peptide we designed for targeting the GREM1–ACVR1C interface effectively disrupts this loop and demonstrates therapeutic potential.

Intercellular communication during embryonic development, tissue repair, and tumor progression is critically orchestrated by secreted signaling molecules, particularly through paracrine and autocrine mechanisms that govern cell fate and behavior(Xie et al., 2020). For instance, factors such as IGF-I and IL-24 coordinate cellular proliferation, differentiation, and microenvironmental remodeling in skeletal development and skin wound healing via both paracrine and autocrine routes(Liu et al., 2023; Wang et al., 2013). Similarly, tumor cells exploit these dual signaling modes to enhance their survival and invasive capacity(Basu et al., 2025). During dissemination to distant sites, metastatic tumor cells have been observed to bring along components of the primary tumor stroma, including CAFs, reflecting their persistent reliance on microenvironmental support(Ao et al., 2015; Hurtado et al., 2020). In the seminal work on TGF, Sporn et al. first proposed that a potential mechanism of malignant transformation involves the autocrine production of growth factors to which the cell itself is responsive, thereby enabling escape from microenvironmental dependence. This mechanism may represent an evolutionary reactivation of autocrine strategies utilized during early embryonic development, allowing cells to maintain survival even in the absence of external support(Sporn and Todaro, 1980; Wydooghe et al., 2017). Weinberg et al. further suggested that paracrine signals from the stroma might trigger the emergence of signaling autonomy in tumor cells. Yet, the exact pathways by which such a shift toward autocrine self-sufficiency is achieved have not been conclusively elucidated(Hanahan and Weinberg, 2000). While neuron-secreted NLGN3 has been reported to upregulate NLGN3 expression in glioma cells, it has not been demonstrated whether this process evolves into a self-sustaining autocrine loop (Venkatesh et al., 2015). Thus, whether tumor cells can transition from paracrine induction to a self-sustaining autocrine circuit, and thereby maintain their malignant phenotype in the absence of the native microenvironment, remains an unresolved question. In this study, we show that during CRC metastasis, GREM1 expression undergoes a stromal-to-epithelial shunt, establishing an autocrine signaling mode initiated by paracrine cues. This transition suggests that metastatic cells acquire signaling autonomy by reactivating developmental autocrine programs, enabling them to survive and disseminate in distant organs with reduced microenvironmental support. Such a shift from “dependence on soil” to “self-construction of soil” reflects the remarkable adaptability of tumor cells and represents a pivotal step in the evolution of metastatic competence.

As a canonical antagonist of BMPs within the TGF-β superfamily, GREM1 plays a pivotal regulatory role across diverse physiological and pathological contexts. During embryogenesis, GREM1 modulates BMP gradients to orchestrate the development of organs such as the kidney(Michos et al., 2004), skeleton(Worthley et al., 2015), and limbs(Malkmus et al., 2021), and is particularly essential for maintaining mesenchymal stemness and tissue integrity in the gut(Worthley et al., 2015). Beyond development, GREM1-mediated BMP signaling is also critically involved in benign conditions including obesity(Grillo et al., 2023), tissue repair(Koppens et al., 2021), and fibrosis(Baboota et al., 2022). In the context of cancer, GREM1 displays marked tissue-specific functionality. In intestinal tumors, studies led by Tomlinson’s group and collaborators revealed that in hereditary mixed polyposis syndrome (HMPS), duplication of an upstream enhancer drives aberrant GREM1 expression in epithelial cells, disrupting the BMP-driven stemness gradient and expanding progenitor-like cells toward the mucosal surface, thereby initiating polyp development(Davis et al., 2015; Jaeger et al., 2012b; Mulholland et al., 2025). GREM1 also reinforces stemness and tumor progression by suppressing BMP signaling in colorectal cancer(Kobayashi et al., 2021). In contrast, in pancreatic ductal adenocarcinoma, GREM1 expression constrains EMT and promotes differentiation toward a less invasive epithelial phenotype(Lan et al., 2022). While traditionally categorized as a BMP antagonist, portraying GREM1 solely as an inhibitory molecule overlooks its diverse and at times paradoxical roles in various biological settings(Gao et al., 2023). Accumulating evidence suggests that GREM1 possesses non-canonical, BMP-independent activities, engaging multiple receptors and activating diverse downstream signaling pathways. Previous studies have implicated EGFR(Park et al., 2020b) and VEGFR2(Mitola et al., 2010) as putative GREM1 targets, though their binding affinities have not been determined. Zhu et al. reported that GREM1 binds to FGFR1 with a dissociation constant of *K*_d_ = 10.6 nM(Cheng et al., 2022). In this study, we further identified ACVR1C(Tsuchida et al., 1996) as a novel binding receptor for GREM1, with a dissociation constant of *K*_d_ = 67.67 ± 10.35 nM. Notably, this binding affinity is markedly higher than that of the canonical ACVR1C ligand Activin B (*K*_d_ = 854.1 ± 127.47 nM), representing a 12.6-fold increase. These results suggest that GREM1 engages ACVR1C with superior affinity, indicating its potential to act as a dominant ligand within this signaling pathway.

Unlike the mechanism described by Lan et al. in pancreatic cancer, where GREM1 inhibits EMT via classical BMP antagonism, our findings demonstrate that GREM1 binds ACVR1C and activates the SMAD2/3 axis independently of BMP signaling. This interaction initiates a positive autocrine feedback loop that promotes EMT and metastasis in CRC. These contrasting roles highlight the context-dependent plasticity of GREM1, which may engage distinct signaling programs across tissue types to drive divergent cellular outcomes. While GREM1 has not been systematically recognized as a member of the cytokine family, emerging evidence suggests it acts as an adipokine in adipose tissue, modulating metabolic homeostasis(Hedjazifar et al., 2020). Our study extends this concept by showing that GREM1 exhibits key cytokine characteristics, including active secretion, receptor engagement, downstream signaling activation, and autocrine feedback amplification. Reframing GREM1 as a cytokine beyond BMP antagonism may provide new insights into its pleiotropic roles in cancer, metabolic disorders, and developmental or regenerative processes.

The rise of cancer genomics and precision medicine has ushered in a new era of molecularly tailored therapies for advanced CRC. Approved targeted agents, including those against VEGF, EGFR, BRAF V600E mutations, and HER2 alterations, have shown efficacy in genetically defined patient subsets(Biller and Schrag, 2021). Yet, challenges such as limited bioavailability, acquired resistance, and narrow indications constrain their widespread utility(Ciardiello et al., 2022). Distinct from mutation-restricted targets, GREM1 is a broadly expressed cytokine with sustained activity across diverse pathological stages. Its tractable biology and context-specific signaling functions highlight GREM1 as an attractive, potentially more universally applicable therapeutic target in CRC. Therapeutic targeting of GREM1 to date has focused largely on full neutralization strategies using monoclonal antibodies. Fully humanized monoclonal antibodies such as Ginisortamab and TST003 have advanced into phase I/II clinical trials(Davies et al., 2023; Rivera et al.; Sarker et al.; Sun et al.). Although these antibodies have shown efficacy in preclinical models, the indispensable roles of GREM1 in physiological processes such as intestinal homeostasis and bone marrow hematopoiesis raise the risk of adverse effects from systemic inhibition, posing a major challenge to their clinical translation(Koppens et al., 2021; Rowan et al., 2020). In recent years, PPIs have evolved from being considered “undruggable” to emerging as attractive therapeutic targets. Among strategies to modulate PPIs, small peptides have demonstrated particular promise due to their high binding affinity and specificity(Lu et al., 2020). In this study, we propose a more selective therapeutic strategy by leveraging the structural interface of the GREM1–ACVR1C interaction. Using structure-guided molecular design, we developed a high-affinity peptide that specifically disrupts this interaction in tumor cells, leading to a marked inhibition of metastatic potential. Unlike conventional neutralizing antibodies that globally suppress GREM1 activity, our approach targets the pathogenic GREM1–ACVR1C axis with precision, offering a highly selective therapeutic avenue for metastatic CRC. This work highlights the translational potential of targeting PPIs as an anticancer strategy through rational molecular intervention.

In summary, our findings reveal that the GREM1–ACVR1C axis acts as a key mediator of the stromal-to-epithelial shunt and the establishment of autonomous GREM1 signaling in CRC. By uncovering this pathway, we not only deepen our understanding of tumor self-sufficiency mechanisms, but also identify a functionally precise intervention point that opens new avenues for disrupting metastatic competence through targeted dismantling of self-reinforcing oncogenic circuits.

## Acknowledgements

We thank Ruohan Li and Changxue Li for assisting with IHC data scoring; Guihua Wang for providing the *APC^Min/+^*mouse line; Pengcheng Bu for assisting with the construction of the cecum to liver metastasis mouse model; Lei Zhou for drawing the GREM1-ACVR1C protein docking; and Xingqiao Xie from the Sample Preparation and Analysis Core Facility of Shenzhen Medical Academy of Research and Translation (SMART) for technical support of MST. This work was supported by grants from the National Natural Science Foundation of China (81874176, 82072766, 32471261), Guangdong Provincial Key Laboratory of Digestive Cancer Research (2021B1212040006), the Sanming Project of Medicine in Shenzhen (SZSM202111005), funding of Shenzhen Clinical Research Center for Gastroenterology (Gastrointestinal Surgery) (LCYSSQ20220823091203008), Guangdong Medical Science and Technology Research Foundation (A2022068).

## AUTHOR CONTRIBUTIONS

H.Z. and N.L. conceived and designed the study and wrote the manuscript. H.Z. performed most of the experiments and analyses. Q.J. and Y.J. helped with RT-qPCR analysis and clinical tissue IF experiments. Z.F. and Y.Y. helped with subcutaneous tumor IF experiments. Y.Y. assisted with the design and drawing of schematics and mechanistic diagrams. Y.G. assisted with histopathological assessment. N.W. and B.Z. performed ACVR1C-ECD and ACVR1C-ECD-doble-mutant proteins expression and purification. J.L. helped with single-cell RNA-seq data analysis. Z.Z. helped with the establishment of the cecum to liver metastasis mouse model. L.G., Y.Z. (Affiliation 1), Y.H., Y.Z. (Affiliation 7), and J.Z. provided intellectual feedback. Note: Y.Z. (Affiliation 1) and Y.Z. (Affiliation 7) are distinct individuals. S.L. and L.F. assisted with data interpretation and analysis and text proofreading. X.W. helped with CHIP experiments. T.W. helped with RNA-seq data analysis. X.S., T.W., Y.J. and N.L. supervised the study.

## METHODS

### Cell lines

Human colon cancer cell lines (HCT116, SW480, DLD1, LOVO, LS174T, and HT29), mouse colon cancer cell line (MC38) and human embryonic kidney cell lines (HEK293 and its derivative 293T) were purchased from ATCC. Colon cancer cells, HEK293 cells and 293T cells were cultured in RPMI1640 (HyClone, SH30809.01) and DMEM (SH30022.01), respectively, supplemented with 10% fetal bovine serum (Biological Industries, 04-001-1ACS) and 1% Penicillin/Streptomycin (Biological Industries, 03-031-1B) at 37°C, 5% CO_2_.

### Patients and tissue samples

A total of 106 archived human colorectal cancer specimens were obtained from the colorectal cancer database and tissue bank of the First Affiliated Hospital of Sun Yat-sen University (SYSU). These tissues were collected from patients who underwent radical resection for colorectal cancer between 2008 and 2015, and followed up until December 2017. All patients had provided written informed consent, and the use of these samples was approved by the Institutional Review Board of the First Affiliated Hospital, SYSU. In addition, a commercial human CRC tissue microarray containing primary tumor tissues from patients with stage I–III CRC (n = 93) was purchased from Shanghai Outdo Biotech Company (Shanghai, China), with ethics approval documented by the company.

### Conditioned medium (CM)

To investigate the effect of secreted GREM1 on CRC cells, an *in vitro* GREM1 secretion system was established using HEK293 cells. Stably GREM1-expressing cells (pcDNA3.1-GREM1) and empty-vector transfected cells (pcDNA3.1) were cultured in DMEM/F12 medium with 10% FBS until 40% confluence. After complete removal of the normal culture medium, HEK293-GREM1 and HEK293-Vec cells were continuously cultured in DMEM/F12 medium without FBS for 5 days before medium collection. GREM1 or Vector conditioned medium (GREM1-CM/Vec-CM) was then centrifuged at 1000 rpm for 30 min and the supernatant was collected for further study.

### Lentiviral plasmid construction, lentivirus production and infection

Human *GREM1* CDS (NM_013372.7) with a HA-mCherry tag or human *ACVR1C* CDS (NM_145259.3) with a Flag-GFP tag was cloned into a pCDH-CMV-MCS-EF1-puro vector. Truncated or point mutations of GREM1 or ACVR1C were cloned from entire GREM1- or ACVR1C-expressing plasmids by PCR. Human *GREM1* CDS (NM_013372.7) was cloned into the pLV-EF1a-IRES-Puro lentiviral vector. CRISPR-mediated gene knockout: The sequences targeting GREM1 were GREM1 KO27 (gRNA1: 5′– GCAAATACCTGAAGCGAGAC –3′) and GREM1 KO28 (g*RNA2:* 5′–AAGCAGACCATCCACGAGGA –3′). The Cas9 lentivirus and gRNA1/2 lentivirus were purchased from GenePharma. shRNA-mediated silencing: The human ACVR1C shRNA target sequences are listed as follows: shACVR1C#1 (5′– CGGAGGAATTGTTGAGGAGTA –3′); shACVR1C#2 (5′– GCAACACCTCAACTCATCTTT –3′). All inserts and vectors were purified from agarose gel using the FastPure ® Gel DNA Extraction Mini Kit (Vazyme, DC301-01) and assembled with Gibson Assembly Master Mix(Gibson et al., 2009) (NEB, E2611) according to the manufacturers’ protocols. All plasmids were verified by Sanger sequencing. HEK 293T cells were seeded at a density expected to reach 70-80% confluence at the time of transfection.

To produce lentivirus, plasmids mentioned above together with packaging plasmid (*psPAX2*) and envelope plasmid (*pMD2.G*) were mixed in a 3.9:2.1:1 ratio and transfected into the cells using polyethylenimine (PEI). After 48-72 hours, supernatant containing lentivirus was collected, filtered, and either used immediately or stored at - 80□ for later applications.

HCT116 and SW480 tumor cells were infected with lentiviral particles in the presence of 5 μg/mL polybrene. To establish stable cell lines, these infected cells were selected with 1.25 μg/mL puromycin for 2 weeks.

### Isolation of fibroblasts from normal and tumoral human intestinal tissues

Fibroblasts were isolated from normal and tumoral human intestinal tissues. Briefly, a cell dissociation buffer was prepared using DMEM supplemented with 10% FBS, 1% Penicillin/Streptomycin, collagenase type D (1 mg/mL, Roche, 11088866001), and DNase I (20 µg/mL, Roche, 10104159001). Tumor tissues were washed twice with DMEM or phosphate-buffered saline (PBS), transferred to a 100-mm culture dish containing 15 mL of cell dissociation buffer, and minced into fragments (< 1 mm³) using sterile razor blades. The tissue fragments were enzymatically digested at 37 °C for 30 min. Following digestion, the cell suspension was filtered through a 70-µm cell strainer to obtain a single-cell suspension. The cell suspension was centrifuged at 1000 rpm for 5 min, and the pellet was resuspended in DMEM. This step was repeated, and the final pellet was resuspended in DMEM supplemented with 10% FBS and 1% penicillin-streptomycin at 37 °C in a humidified incubator with 5% CO□.

### Animal experiments

The immunocompromised nude(Flanagan, 1966) and NOG(Ito et al., 2002) female mice (6 weeks old) were purchased from Guangzhou Vital River Laboratory Animal Technology Co., Ltd. *Grem1-CreER^T2^*(stock no. 027039)(Worthley et al., 2015), *Rosa-mTmG* (stock no. 007576)(Muzumdar et al., 2007), *Rosa-LSL-DTA* (stock no. 007900) mice(Voehringer et al., 2008) were obtained from the Jackson Laboratory. *APC^Min/+^*mice were obtained from the Gempharmatech Co., Ltd (stock no. 002020)(Moser et al., 1990). *Grem1-CreER^T2^*mice were crossed with *Rosa-mTmG* mice or *Rosa-LSL-DTA* mice to generate GR or GD mice, respectively. The Cre recombinase activity was induced by the ER antagonist tamoxifen (TMX), allowing Grem1^+^ cells to express GFP in GD mice. GD mice were further crossed with *APC^Min/+^* mice to generate AGD mice. AGD and control mice were administered with 100mg/kg tamoxifen (TMX) through oral gavage at 4-week-old time, when tumor initiates. In AGD and its control mice, activation of Cre lead to the expression of DTA (diphtheria toxin A chain), which removed the population of Grem1^+^ cells from the *APC^Min/+^* mice. All animals were maintained at the Animal Experiment Center of Sun-Yat-Sen University, and all procedures were approved by the Animal Care and Use Committee of Sun-Yat-Sen University. Mice were randomized at the beginning of each experiment. For tail vein-to-lung metastasis model, 5×10^5^ MC38-luc cells were resuspended in 100 μl of PBS and injected into the tail veins of GD mice or *Grem1-CreER^T2^/Rosa-LSL-DTA* mice (n = 5 mice in each group); 1×10^6^ HCT116-luc cells, transduced with lentivirus carrying a control shRNA or two ACVR1C shRNAs or carrying a pLV or pLV-GREM1, were injected into the tail veins of nude mice (n = 5 mice in each group). For spleen-to-liver metastasis model, 5×10^5^ MC38-luc cells were resuspended in 50 μl of PBS and injected into the spleen of GD mice or *Grem1-CreER^T2^/Rosa-LSL-DTA* mice (n = 5 mice in each group). 5×10^5^ HCT116-luc cells carrying a pLV or pLV-GREM1 were resuspended in 50 μl of PBS and intrasplenically injected in NOG mice (n = 5 mice in each group). ACVR1C peptide was administered via tail vein injection at a dose of 10 mg/kg once every other day. For cecum-to-liver metastasis model, 1×10^6^ MC38-luc cells were resuspended in 50 μl of PBS and injected into the cecum of GD mice or *Grem1-CreER^T2^/Rosa-LSL-DTA* mice (n = 6 mice in each group); 5×10^6^ HCT116-luc cells transduced with or without lentivirus carrying a pLV or pLV-GREM1 were injected into the cecum of NOG mice (n = 5 mice in each group). SB505124 was administered via intraperitoneal injection at a dose of 10 mg/kg once every other day. The metastases were examined every 5 days post injection using an IVIS Lumina Imaging System. Mice were euthanized between 2-6 weeks after injection. For subcutaneous transplantation, 1×10^6^ HCT116 cells, either unmodified or transduced with lentivirus carrying a control shRNA, two ACVR1C shRNAs, pLV, or pLV-GREM1, were subcutaneously injected into mice (n = 6-8 mice in each group). Mice were euthanized 4 weeks after injection. The tumor tissues were collected for further evaluation.

### Immunohistochemical (IHC) staining

Immunohistochemical staining of GREM1 (1:50, Biorbyt, orb10741), ACVR1C (1:50, Thermo, PA587475) and Ki67 (1:100, Servicebio, GB111499) was performed on primary tumors tissues. After dewaxing, hydration, and antigen retrieval, the rest of the experimental procedures were performed according to the instructions of the SP Immunohistochemistry Kit (ZSBIO, PV9000). Finally, after DAB staining, hematoxylin re-staining, and neutral resin sealing, the sections were observed under a microscope. Images were taken with a Slide Scanning Imaging System (SQS-1000, sqray). Quantification of positive staining was performed using Fiji (ImageJ).

### Immunofluorescence (IF) staining

Tissue was fixed in 4% paraformaldehyde (Thermo Scientific, I28800) for 24 h at 4□°C, washed with PBS, embedded in paraffin, and sectioned at 5 μm thickness. Antigen retrieval was performed using target retrieval solution, pH 9.0 in a pressure cooker for 15–20 min. Non-specific binding was then blocked with 10% normal donkey serum (Abcam, ab7475) and 0.3% Triton X-100 in PBS for 30 min at room temperature. Cells for IF were fixed with 4% paraformaldehyde for 20 min at room temperature, washed with PBS, and permeabilized with 0.2% Triton X-100 in PBS for 20 min. Cells were then blocked in PBS with 5% BSA for 30 min at room temperature. Subsequently, the samples were incubated with goat anti-GREM1 (3 µg/mL, R&D, AF956), mouse anti-β-Catenin (1:100, BD, 610154), rabbit anti-β-Catenin (1:100, Absea, RC-6352), rabbit anti-Vimentin (1:100, CST, 5741), rabbit anti-CD68 (1:100, CST, 26042), rabbit anti-FAP (1:50, Proteintech, 15384-1-AP), rabbit anti-α-SMA (1:100, Abcam, ab5694), rabbit anti-E-Cadherin (1:200, CST, 3195), rabbit anti-ACVR1C (1:50, Thermo, PA587475), rabbit anti-Snail (1:200, Abcam, ab224731) overnight at 4□°C. The tissues were incubated with Alexa-Fluor-conjugated secondary antibodies (Invitrogen) in PBS with 1 % normal donkey serum for 1 h at room temperature. DAPI was then used for counterstaining the nuclei, and images were obtained by a laser scanning confocal microscope (LSM880, Zeiss).

### Analysis of scRNA-seq data

Single-cell RNA sequencing (scRNA-seq) data from the Gene Expression Omnibus (GEO) database colorectal cancer datasets (GSE200997 and GSE221575) were processed using the R ‘Seurat’ package (v4.4). Initial quality control involved rigorous filtering of low-quality cells: Cells expressing fewer than 200 genes or more than 10,000 genes were excluded, and cells with mitochondrial gene content exceeding 25% were discarded to remove potential apoptotic cells or debris. After quality control, a total of 34,675 high-quality cells were retained for downstream analysis. Gene expression matrices were normalized using the “LogNormalize” method implemented in the NormalizeData function, which scales feature counts per cell by total expression and multiplies by a scale factor (10,000), followed by natural log transformation. To identify biologically relevant features, the FindVariableFeatures function was employed to select the top 2,000 highly variable genes (HVGs) exhibiting the highest cell-to-cell variation. Dimensionality reduction was performed using principal component analysis (PCA) on scaled expression data of the identified HVGs. To address technical batch effects between samples and datasets, we applied multiple Canonical Correlation Analysis (CCA) as implemented in Seurat’s integration workflow. Cell clustering was performed using a graph-based approach: The FindNeighbors function constructed a shared nearest neighbor (SNN) graph based on the first 30 principal components, followed by the FindClusters function using the Louvain algorithm at a resolution of 0.8 to identify distinct cell subpopulations. Finally, non-linear dimensionality reduction was achieved through t-distributed Stochastic Neighbor Embedding (t-SNE) using the same principal components.

### Immunoprecipitation (IP)

HCT116 cells were transfected with the indicated plasmids and lysed in NP40 lysis buffer (Beyotime, P0013F) supplemented with protease inhibitor cocktail (Thermo, 78446). Lysates were incubated with the indicated Anti-Flag nanobody magarose beads (Ktsm-life, KTSM1338), Anti-HA nanobody magarose beads (Ktsm-life, KTSM1335) or Anti-GFP nanobody magarose beads (Ktsm-life, KTSM1334) overnight at 4□°C. The protein complex was washed four times with the NP40 lysis buffer, eluted with 1×loading buffer (Beyotime, P0015) by boiling for 5 min, followed by mass spectrometry and immunoblotting with the indicated antibodies.

### Mass spectrometry (MS) analysis

Proteins were separated by 10% SDS-PAGE and visualized using Coomassie Brilliant Blue staining before mass spectrometry analysis. The stained gel bands were excised (∼1–2 mm), washed with MilliQ water, and destained using 25 mM NH□HCO□ and 50% acetonitrile (ACN) at 37°C. The gel pieces were dehydrated with ACN, reduced with 10 mM dithiothreitol (DTT) in 25 mM NH□HCO□ at 37°C for 1 h, and alkylated with 30 mM iodoacetamide (IAA) in 25 mM NH□HCO□ in the dark for 45 min. After sequential washing with MilliQ water and 50% ACN, the gel pieces were dehydrated with ACN and digested overnight at 37°C with trypsin (20 ng/μL) in 25 mM NH□HCO□. Peptides were extracted using 60% ACN followed by pure ACN, pooled, lyophilized, resuspended in 0.1% formic acid (FA), and purified using ZipTip C18 before analysis. Mass spectrometry was performed using a Thermo Fisher Orbitrap HF-X coupled with an Easy-nLC 1200 system and a C18 column, employing a 90-min gradient of 5–35% ACN in 0.1% FA at a flow rate of 300 nL/min. MS1 scans were acquired at a resolution of 60,000 with an AGC target of 3 × 10□, a maximum injection time of 20 ms, and a scan range of m/z 350–1800. MS2 scans were performed at a resolution of 15,000 with an AGC target of 2 × 10□, a maximum injection time of 100 ms, TopN of 20, and a normalized collision energy (NCE) of 32. Raw MS data were analyzed using Proteome Discoverer 2.4, with protein identification performed against the SwissProt human database using trypsin specificity (allowing one missed cleavage site), cysteine alkylation with MMTS, a precursor mass tolerance of 10 ppm, a fragment mass tolerance of 0.02 Da, and a false discovery rate (FDR) threshold of <1%.

### Immunoblotting (IB)

Protein was extracted from the cells with RIPA buffer (Beyotime, P0018) or NP40 lysis buffer (Beyotime, P0013F) and separated by SDS-PAGE, and transferred to polyvinylidene difluoride membranes. Primary antibodies against GREM1 (1:1,000, SinoBiological, 50016-R117), ACVR1C (1:1,000, Thermo, PA587475), Flag-tag (1:1,000, CST, 14793), HA-tag (1:1,000, CST, 3724), E-Cadherin (1:1,000, CST, 3195), β-Catenin (1:1,000, CST, 8480), ZEB1 (1:1,000, CST, 3396), Snail (1:1,000, CST, 3879), SMAD2/3 (1:1,000, CST, 8685), p-SMAD2/3 (1:1,000, CST, 8828), SMAD1 (1:1,000, CST, 6944), p-SMAD1/5/9 (1:1,000, CST, 13820), TGFβ (1:1,000, CST, 3709), TGFβR1 (0.3µg/mL, R&D, AF3025), β-actin (1:5,000, Beyotime, AF0003) and GAPDH (1:5,000, Beyotime, AF0006) were used in this study. Peroxidase-conjugated secondary antibody (1:10,000, Cell Signaling Technology, 7074, 7076) was used and signal was visualized using an enhanced chemiluminescence assay (ECL, Thermo), according to the manufacturer’s protocol. Band intensity was quantified using Fiji (ImageJ) by grayscale analysis.

### Recombinant protein production and purification

Expi293F cells were transfected with a *pcDNA3.4-ACVR1C-ECD-Fc* and *pcDNA3.4-ACVR1C-ECD-Fc-double mutant (E85A/T101A)* expression vector to produce the target protein, which was subsequently purified using a Protein G column. Briefly, the coding sequence (CDS) of the human *ACVR1C* extracellular domain (*ACVR1C-ECD*, NM_145259.3, residues 1-339) fused to an Fc tag was cloned into the pcDNA3.4 vector. Expi293F cells were transfected with this construct, the supernatant was harvested 5 days post transfection.

The supernatant was first centrifuged at 1000 rpm for 20 minutes to remove cell debris, and the supernatant was further centrifuged at 8000 rpm for 30 minutes, followed by filtering with a 0.45 μm PES filter. The protein in the supernatant was then purified using a Protein G column equilibrated with binding buffer (0.15 M NaCl, 20 mM Na□HPO□, pH 7.0). The target protein was eluted with 0.1 M glycine (pH 2.5) and immediately neutralized with 1 M Tris-HCl (pH 8.5).

Subsequently, the protein buffer was exchanged into a 20 mM Tris-HCl (pH 7.5) system. To further purify the sample, it was centrifuged at 12000 rpm and 4°C for 10 minutes to remove impurities and precipitates. The clarified sample was then loaded onto an ion exchange column (HiTrap™ Capto™ Q ImpRes) equilibrated with binding buffer (20 mM Tris-HCl, pH 7.5). The ACVR1C-ECD-Fc protein, having an opposite charge to the resin, was bound to the column. Finally, the target protein was eluted with a linear gradient (0-100%) of elution buffer (20 mM Tris-HCl, 1 M NaCl, pH 7.5) over 6 column volumes.

### Protein pull-down assay

Protein pull-down assay was performed using purified recombinant human His-tagged GREM1 protein and recombinant human ACVR1C-ECD and Fc chimera protein. Protein was enriched by Pierce Protein A magnetic beads (MCE, HY-K0202) or Ni Sepharose 6 Fast Flow (GE, 17531801) following the manufacturer’s instructions. Pulled-down proteins were detected by Coomassie Brilliant Blue staining.

### MicroScale thermophoresis (MST)

MST was carried out on a Monolith NT.115 instrument (NanoTemper Technologies GmbH). To evaluate ACVR1C-ECD or ACVR1C peptide or ACVR1C-ECD-double mutant (E85A/T101A) binding to GREM1-His or ACTIVINB-His, an increasing concentration of purified ACVR1C-ECD-Fc protein (0–27.5 μM) or ACVR1C peptide (0–2.3 μM) or ACVR1C-ECD-double mutant (0–27.5 μM) was incubated with 50 nM RED-labeled (NanoTemper Technologies GmbH) GREM1-His protein (R&D, 5190-GR) or ACTIVINB-His protein (SinoBiological, 10814-H08H). Experiments were carried out in a PBS buffer pH 7.4 using premium capillaries.

### Protein-protein interaction docking study

GREM1 (PDB: 5AEJ) was selected as the ligand and ACVR1C (PDB: AF-Q8NER5-F1) as the receptor for protein-protein docking. The HDOCK web service was used for docking with default parameters (http://hdock.phys.hust.edu.cn/). Key amino acid residues in the binding pocket between GREM1 and ACVR1C were further identified based on the docking module(Yan et al., 2017).

### RNA-seq and gene set enrichment analysis (GSEA)

Total RNA was extracted using Trizol reagent (Invitrogen, 15596026) and quantified with a NanoDrop spectrophotometer (Thermo Fisher Scientific). RNA integrity was assessed using an Agilent 2100 Bioanalyzer. mRNA was enriched using oligo(dT) magnetic beads, fragmented, and reverse-transcribed into cDNA. After adapter ligation and PCR amplification, libraries were sequenced on an Illumina platform, generating 150-bp paired-end reads. Raw reads were trimmed and aligned to the human reference genome (GRCh38) using STAR. Differential gene expression analysis was conducted using Limma, with significance thresholds set at |log2FoldChange| >1.5 and adjusted P-value <0.05. Gene set enrichment analysis (GSEA) was performed using the GSEA software (Broad Institute) with the MSigDB gene sets to identify enriched biological pathways, employing 1,000 permutations and FDR <0.25 as the cutoff for significance.

### RT–qPCR

Total RNA was extracted using Trizol reagent (Invitrogen, 15596026). According to the instruction, cDNA was generated using the PrimeScript RT reagent Kit with gDNA Eraser (Accurate Biology, AG11706). The SYBR Green Premix Pro Taq HS qPCR Kit (Accurate Biology, AG11701) was then used to quantify mRNA expression according to the manufacturer’s instruction. All results were calculated using the 2^−ΔΔct^ method. Primers used in the study are listed in Supplementary Table 1.

### ChIP

SW480 cells were starved in DMEM with 1% FCS overnight before treatment with vehicle, 10 μM SB505124 for 24 hours. Cells were fixed in 1 % paraformaldehyde for 10 min at RT for DNA-protein cross-linking, followed by quenching with glycine. Cross-linking chromatin was prepared using the SimpleChIP® Enzymatic Chromatin IP Kit (CST, 9002) according to the manufacturer’s instructions. For immunoprecipitation, 10 μg chromatin was incubated with 10 μL anti-histone H3 rabbit IgG (CST, 14269, positive control), 2 μL normal Rabbit IgG (CST, 2729) or 5 μL anti-SMAD2/3 rabbit IgG (CST, 8685) at 4 °C overnight. 2% chromatin prior to immunoprecipitation was used as input. Chromatin-protein-antibody complex was captured by protein G magnetic beads, and chromatin was released by reversal of cross-links and purified using the SimpleChIP® Enzymatic Chromatin IP Kit (CST, 9002) according to the manufacturer’s instructions. DNA was quantified by qPCR with primers targeting predicted SMAD2/3/4 binding regions on *GREM1* or *SNAI1* promoters. DNA levels were normalised to the input, and the fold-change of enrichment was calculated over the control. ChIP–qPCR primers are listed in Supplementary Table 2.

### Scratch assay

Cells were seeded into 6-well plates after centrifugation and digestion with 0.05% trypsin. When the cell density reached 90%, three vertical lines were scratched in each well with a 10 μL pipette tip and the floating cells were gently washed away with 1×PBS. Complete medium was added, and images of the scratch area were taken at 0 h. Three different fields of view were selected for each well. After photography, the medium was replaced with serum-free medium. Wound healing was documented at the same location after 24 h or 48 h of incubation.

### Transwell invasion assay

Cells (1×10^5^) were seeded in serum-free medium in the Matrigel-coated (Corning, 354480) transwell chambers (24-well insert, 8-μm pore size; BD Biosciences) for invasion experiments. The lower chamber was filled with RPMI1640 or DMEM containing 20% FBS. The migration of HCT116 and SW480 cells was measured in three random visual fields and quantified by microscopy after 48 h of incubation, followed by staining with DAPI or crystal violet. The invasive capacity of the cells was assessed using ImageJ software for quantification.

### Statistical analysis

All the statistical analyses were performed using GraphPad Prism 9, and error bars indicate s.e.m. Student’s t-test assuming equal variance and one-way analysis of variance for independent variance were used. Growth curves were generated using ANOVA for repeated measurement. P<0.05 was considered significant. The number of independent experiments, the number of events and information about the statistical details and methods are indicated in the relevant figure legends.

