## Supplementary Materials for "A Paracrine-to-Autocrine Shunt of GREM1 Fuels Colorectal Cancer Metastasis via ACVR1C"

^14^ Lead contact

* Corresponding author:

Ningning Li

Xuetong Shen

**SUPPLEMENTAL FIGURE LEGENDS**

**Figure S1.** **The stromal GREM1^+^ cells are CAFs along the intestinal isthmus** **and are distinct from the myofibroblast population.** (A) Schematic of the GR (*Grem1-CreER^T2^;Rosa-mTmG*) mouse model. The black triangle indicates the loxP sites. (B) Representative images of haematoxylin and eosin (H&E) staining (Left) and IHC staining of GFP (Right) in GR mice. Mice received TMX (100 mg/kg) by oral gavage for five consecutive days starting at 24 weeks of age, followed by analysis. Scale bars, 100 μm and 25 μm. (C) Representative images for IF staining of GFP, E-cad and α-SMA in GR mice. (D) Representative images of GREM1 IHC staining on human primary CRC-adjacent normal tissue. (E–I) Representative images for IF staining of GREM1 and β-CAT (E), VIM (F), CD68 (G), FAP (H), α-SMA(I) in human stage I-III CRC tumors. Scale bars, 20 μm. (J) Quantification of the number of GREM1^+^ cells among the populations of CRC cells (β-CAT^+^), stromal cells (VIM^+^), macrophage cells (CD68^+^), activated fibroblasts (FAP^+^) and myofibroblasts (α-SMA^+^) (n = 5 patients). The bar charts show the mean ± s.e.m.

**Figure S2. Expression patterns of GREM1 in single-cell transcriptome of CRC tissues.** (A) t-SNE visualization of single-cell clustering showing major cell types in CRC tissues. Individual cells represented by dots colored by annotated cell type. (B) GREM1 expression levels across the single-cell landscape. Expression intensity is shown by color gradient (low: light purple; high: dark red). (C) GREM1 expression distribution across cell types. (D) GREM1 expression in fibroblasts across CRC stages. (E) GREM1 expression in epithelial cells across CRC stages. All statistical comparisons were performed using non-parametric Wilcoxon tests. t-SNE: t-distributed stochastic neighbor embedding.

**Figure S3. High ACVR1C expression marks advanced CRC and poor outcome.** (A) MS analysis identified ACVR1C as the cytoplasmic membrane protein from HCT116 cells pulled down by GREM1-his. Four peptide fragments were detected by MS that matched the ACVR1C protein sequence, supporting its identification. (B) Increasing concentrations of recombinant ACVR1C-ECD double mutant protein (0-27.5 μM) were incubated with red-labeled 50 nM recombinant GREM1-his protein. MST was used to evaluate ACVR1C double mutant binding to GREM1-his. (C) Representative images of ACVR1C IHC staining in human primary CRC tissues from SYSU and a commercial tissue microarray. (D) Quantification of epithelial ACVR1C expression across stage I–IV primary CRC tissues based on IHC staining shown in c. Data are mean ± s.e.m. *P* values were calculated using one-way ANOVA with Bonferroni multiple-comparison test. (E) ScRNA-seq analysis of ACVR1C expression in epithelial cells across CRC stages. Statistical comparisons were performed using non-parametric Wilcoxon tests. (F) Kaplan-Meier survival curves for stage IV CRC patients from SYSU, stratified by epithelial ACVR1C expression levels. Patients were divided into low and high expression groups based on the median expression level. Significance was determined using a two-sided log-rank test. HR, hazard ratio.

**Figure S4. GREM1-CM induces EMT****,** **migration and invasion of CRC cells via the TGFβ superfamily signaling pathway.** (A) Schematic diagram of the GREM1 secretion system established in HEK293 cells. The GREM1 expression plasmid was stably transfected into HEK293 cells, and empty vector-transfected HEK293 cells served as controls. 5 days after culturing, the culture medium (supernatant) and cell pellets were collected separately; the supernatant was used as the GREM1 conditioned medium (GREM1-CM) or control conditioned medium (Vec-CM). (B) RT–PCR analysis confirmed the expression of *GREM1* in GREM1-transfected HEK293 cells. *ACTB* was used as a loading control. (C) Immunoblotting analysis showed that GREM1 protein was detected in conditioned medium of HEK293 cells. (D, E) Quantification of mRNA levels of epithelial marker *CDH1* and mesenchymal markers *CTNNB1*, *SNAI1*, and *ZEB1* by RT–qPCR in HCT116 (D) and SW480 (E) cells treated with Vec-CM, GREM1-CM, GREM1-CM + GREM1 BAb, or GREM1-CM + IgG. *ns*: not significant. (F) Representative images of scratch assays (Upper) and Transwell invasion assays (Lower) for both HCT116 and SW480 cells treated with Vec-CM, GREM1-CM, GREM1-CM + GREM1 BAb, or GREM1-CM + IgG. (G) Quantification of the relative migration rate in HCT116 cells at 24 hours, showing that GREM1-CM treatment significantly increased migration compared to Vec-CM, and that GREM1 BAb inhibited this effect. (H) Quantification of the relative migration rate in SW480 cells at 48 hours, showing a similar trend as in HCT116 cells. (I) Quantification of the invasion rate from Transwell assays for HCT116 and SW480 cells treated with the indicated conditions. The invasion rate was significantly reduced by GREM1 BAb in both cell lines. (J) GSEA was performed using the KEGG gene set on the significantly differentially expressed genes in HEK293 cells treated with GREM1-CM or Vec-CM. The analysis indicated that the TGFβ signaling pathway was the most significantly enriched. For D, E, and G–I, n = 3 independent experiments. Columns represent the mean of triplicate experiments. Data are presented as mean ± s.e.m. *P* values were calculated using one-way ANOVA with Bonferroni multiple-comparison test. Scale bars, 100 μm.

**Figure S5. Knockdown of ACVR1C inhibits the GREM1-CM-induced EMT, migration and invasion in CRC.** (A, B) Expression levels of the epithelial marker *CDH1* and mesenchymal markers *ZEB1*, *SNAI1* and *CTNNB1* were analyzed by RT–qPCR in SW480 (A) and HCT116 (B) cells transfected with Scramble shRNA or shACVR1C and treated with Vec-CM or GREM1-CM. (C) The impact of GREM1-CM on cell migration and invasion following ACVR1C knockdown was evaluated in SW480 and HCT116 CRC cells. Left: Scratch assays were performed using cells transduced with Scramble shRNA or shACVR1C, and cultured in Vec-CM or GREM1-CM. Wound closure was monitored at 0 and 48 hours. Right: Transwell invasion assays were conducted under the same conditions and cells that invaded through Matrigel-coated membranes were stained by crystal violet. (D-G) Quantification of migration (D, E) and invasion (F, G) rates under the indicated conditions. For A, B and D-G, n = 3 independent experiments. Data are presented as mean ± s.e.m. *P* values were calculated using one-way ANOVA with Bonferroni multiple-comparison test. *ns*: not significant. Scale bars, 100 μm.

**Figure S6. p-SMAD2/3 disruption inhibits GREM1-CM-induced EMT, migration and invasion in CRC.** (A, B) Expression levels of the epithelial marker *CDH1* and mesenchymal markers *CTNNB1* and *SNAI1* were quantified by RT–qPCR in HCT116 (A) and SW480 (B) cells cultured with Vec-CM, GREM1-CM, or GREM1-CM in the presence of the SMAD2/3 phosphorylation inhibitor SB505124 (GREM1-CM + SB505124). (C) Representative images of scratch assays (Upper) and Transwell invasion assays (Lower) for both HCT116 and SW480 cells treated with Vec-CM, GREM1-CM or GREM1-CM + SB505124. Scratch closure was monitored at the indicated time points; invasive cells in Transwell assays were stained by DAPI. (D, E) Quantification of relative migration rates in HCT116 (D, 24 h) and SW480 (E, 48 h) cells treated with Vec-CM, GREM1-CM, or GREM1-CM + SB505124. GREM1-CM significantly enhanced migration in both cell lines, an effect that was suppressed by SB505124. (F) Quantification of the invasion rate from Transwell assays for HCT116 and SW480 cells treated with the indicated conditions. The invasion rate was significantly reduced by SB505124 in both cell lines. Columns represent the mean of triplicate experiments. (G) Predicted SMAD2/3/4 binding motifs within the *SNAI1* promoter region, identified using the JASPAR database. Red boxes indicate putative binding sites located upstream of the TSS. (H) ChIP-qPCR analysis of SMAD2/3 occupancy at predicted binding sites (-967, -787, -186 bp) in the *SNAI1* promoter in SW480 cells under the indicated treatment conditions. Chromatin was immunoprecipitated using anti-SMAD2/3 or control IgG, and enrichment of target DNA was quantified by qPCR. DNA levels were normalized to input and expressed as fold change relative to the IgG control. *ns*: not significant. (I) Schematic diagram illustrating the proposed molecular pathway and experimental interventions demonstrating how GREM1-mediated signaling promotes EMT and metastatic potential. For A, B, D-F and H, n = 3 independent experiments. Data are presented as mean ± s.e.m. *P* values were calculated using one-way ANOVA with Bonferroni multiple-comparison test. Scale bars, 100 μm.

**Figure S7.** **Depletion of GREM1^+^ cell impedes the metastasis of CRC in a variety of metastatic models.** (A, B) Number (A) and size (B) of tumors per mouse in AGD models and control (n = 5 mice per group). (C, D) Expression of *Grem1* in intestinal tissue of GD and control mice was confirmed by RT–qPCR (C, n = 3 mice per group) and immunoblotting analysis (D). (E) Schematic representation of the tail vein-to-lung metastasis model. (F) Schematic representation of the spleen-to-liver metastasis model. (G) Schematic representation of the cecum-to-liver metastasis model. (H) Representative images of GD model mice and control mice intravenously injected with MC38-luc cells. Lung metastases were imaged and quantified by IVIS Lumina Imaging System (n = 5 mice per group). (I) Representative images of GD model mice and control mice intrasplenically injected with MC38-luc cells. Liver metastases were imaged and quantified by IVIS Lumina Imaging System (n = 5 mice per group). (J, K) Representative images of the colorectum, and liver from GD model mice and control mice injected into the cecum wall with MC38-luc cells. Colorectal tumor (J) and liver metastases (K) were imaged and quantified by IVIS Lumina Imaging System (n = 5 mice per group). For A-C and J, data are presented as mean ± s.e.m. *P* values were calculated using two-tailed Student’s t-test. *ns*: not significant.

**Figure S8. Knockdown of ACVR1C abolishes GREM1-CM-induced EMT and metastasis in CRC *in vivo*.** (A-C) Images (A), tumor weight (B), and tumor volume (C) of subcutaneous tumors from three groups of nude mice (n = 8 per group). Mice were subcutaneously injected with HCT116 cells transduced with either Scramble shRNA or shACVR1C#1 and shACVR1C#2, all pretreated with GREM1-CM. (D) RT–qPCR analysis of mRNA levels of epithelial marker *CDH1* and mesenchymal markers *CTNNB1*, *SNAI1* and *ZEB1* in subcutaneous tumors. n = 3 independent experiments. (E-G) Representative IF images (E) and quantification (F, G) of E-CAD (green) and SNAIL (red) expression in subcutaneous tumors. Knockdown of ACVR1C reduced SNAIL expression and increased E-CAD expression. Fluorescence intensity was quantified in 3-5 mice per group. (H) Representative IVIS images of lung metastases in nude mice intravenously injected via tail vein with HCT116-luc cells transduced with Scramble shRNA or shACVR1C, all pretreated with GREM1-CM. Luminescence signals from metastatic lesions were monitored over time using the IVIS Lumina Imaging System. Fluorescence signals in the harvested lungs were visualized at day 21 post-tumor injection (upper right panel). For B, C, D, F and G, data are presented as mean ± s.e.m. *P* values were calculated using one-way ANOVA with Bonferroni multiple-comparison test.

**Figure S9.** **SB505124 inhibits GREM1-CM-induced EMT and metastasis of CRC *in vivo*.** (A-C) Images (A), tumor weight (B), and tumor volume (C) from nude mice (n = 6 mice per group) subcutaneously injected with HCT116 cells that were pretreated with Vec-CM, GREM1-CM, or GREM1-CM followed by intraperitoneal administration of the inhibitor SB505124 (10 mg/kg, every other day; GREM1-CM + SB505124). (D) RT–qPCR analysis of epithelial marker *CDH1* and mesenchymal markers *CTNNB1, SNAI1, ZEB1* in subcutaneous tumor tissues from the indicated treatment groups. n = 3 independent experiments. (E-G) Representative IF images (E) and quantification (F, G) of E-CAD (green), SNAIL (red), and DAPI (blue) staining in tumors from the indicated groups. SB505124 treatment restored E-CAD expression and suppressed SNAIL expression (n = 5 mice per group). (H) Representative liver images of NOG mice that were injected with HCT116-luc cells in the cecum wall and treated with either SB505124 or vehicle via abdominal injection. Liver metastases were imaged and quantified using the IVIS Lumina Imaging System (n = 5 mice per group). For B, C, D, F and G, data are presented as mean ± s.e.m. *P* values were calculated using one-way ANOVA with Bonferroni multiple-comparison test.

**Figure S10.** **Autocrine GREM1 and GREM1-ACVR1C binding promote EMT and metastasis of CRC *in vivo*.** (A-C) Images (A), tumor weight (B), and tumor volume (C) from nude mice (n = 6 mice per group) subcutaneously injected with HCT116 cells stably infected with control lentivirus (pLV) or GREM1-overexpressing lentivirus (pLV-GREM1). Autocrine GREM1 significantly increased tumor weight and volume. (D) mRNA levels of epithelial marker *CDH1* and mesenchymal markers *VIM*, *SNAI1* and *ZEB1* were compared by RT–qPCR analysis in subcutaneous tumors derived from HCT116 cells infected with pLV or pLV-GREM1 lentiviruses. (E-G) Representative IF images (E) and quantification (F, G) of E-CAD (green), SNAIL (red), and DAPI (blue) staining in subcutaneous tumors formed by HCT116-luc cells infected with pLV or pLV-GREM1 lentiviruses. pLV-GREM1 reduced E-CAD expression and increased SNAIL expression (n = 5 mice per group). (H) Representative images of Ki67 IHC staining in the lungs of mice intravenously injected with HCT116 cells infected with pLV or pLV-GREM1 lentivirus. Lung metastases larger than 0.1 mm were identified and quantified based on Ki67-positive regions (n = 5 mice per group). (I) Representative IVIS images of liver metastases in NOG mice orthotopically injected in the cecum wall with HCT116-luc cells infected with pLV or pLV-GREM1 lentiviruses. Luminescence signals from liver metastases were imaged and quantified using the IVIS Lumina Imaging System (n = 5 mice per group). (J) Increasing concentrations of recombinant ACVR1C peptide (0–2.3 μM) were incubated with red-labeled 50 nM recombinant GREM1-his. MST was used to evaluate ACVR1C peptide binding to GREM1-his (n = 3 independent experiments). (K) Representative IVIS images of liver metastases in NOG mice (n = 5 mice per group) injected in the spleen with HCT116-luc cells infected with pLV, pLV-GREM1 or pLV-GREM1 combined with ACVR1C peptide treatment. The ACVR1C peptide (10 mg/kg) was administered intravenously every other day, starting 48 hours after cell injection. Luminescence signals were recorded at days 0, 7 and 15 using the IVIS Lumina Imaging System. For B-D, F, G, and J, data are presented as mean ± s.e.m. *P* values were calculated using two-tailed Student’s t-test.

**Figure S11.** **A model for paracrine-driven autocrine of GREM1 boosts metastatic potential of CRC cells.**

**Figure 1**

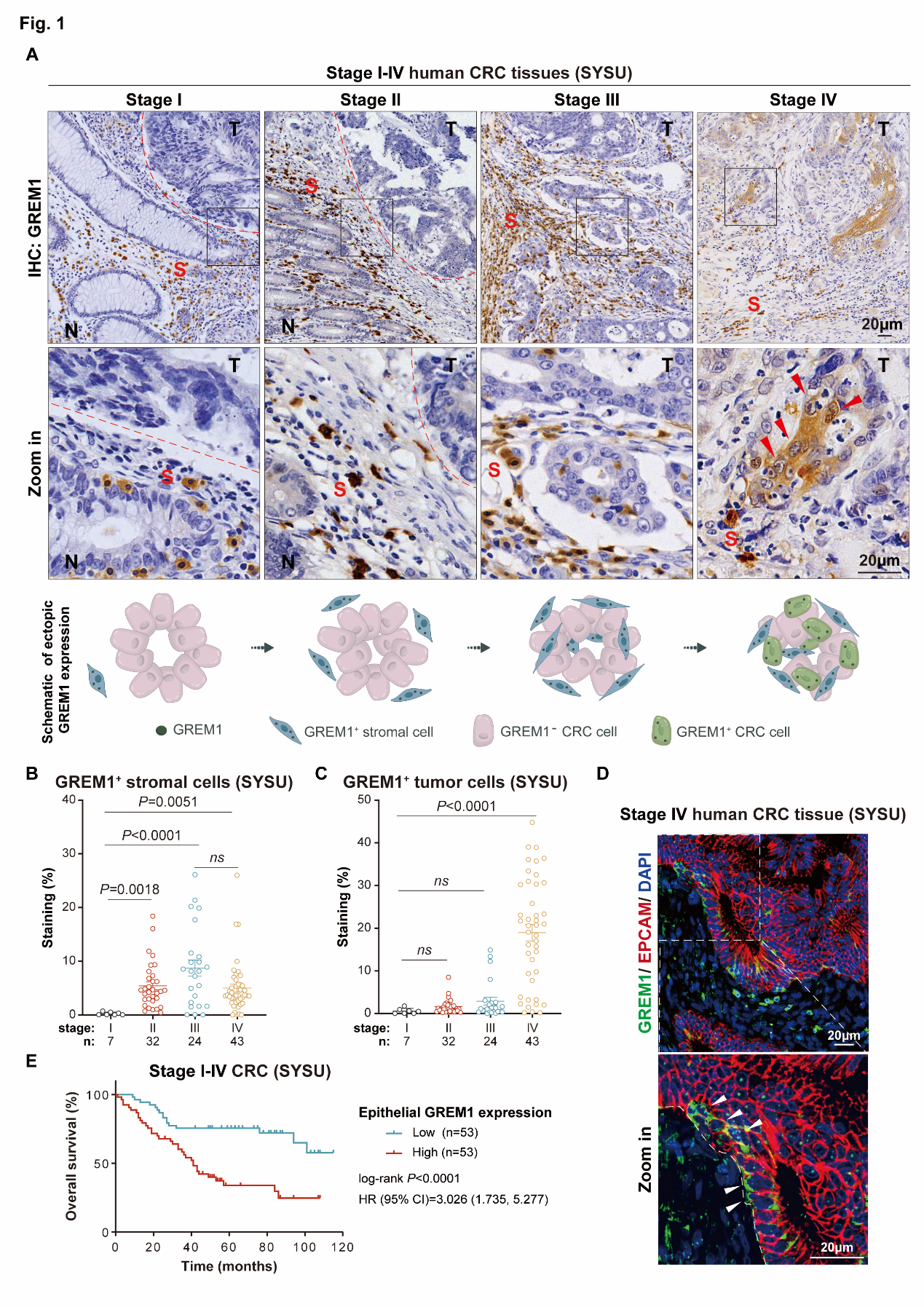

**Figure 2**

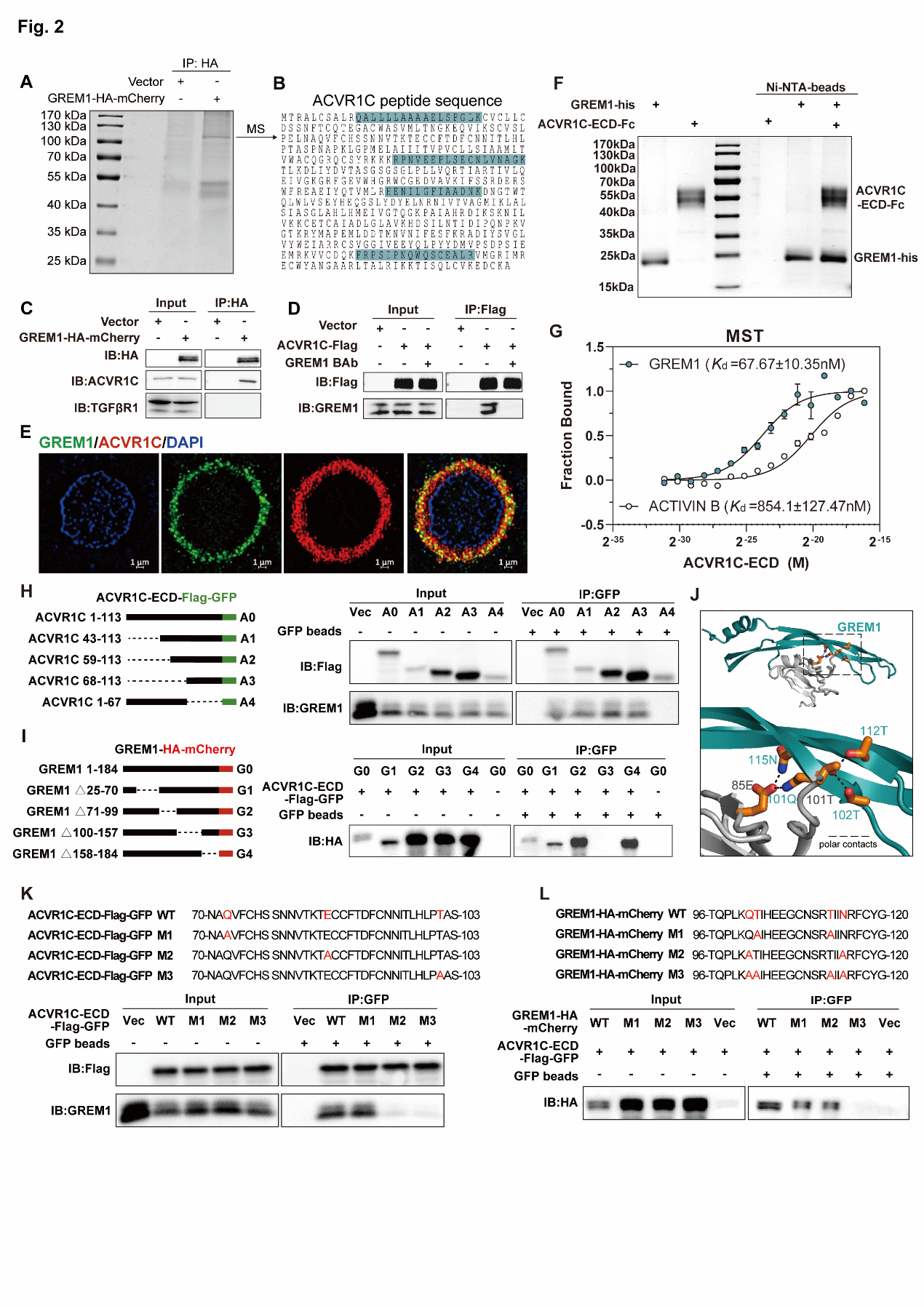

**Figure 3**

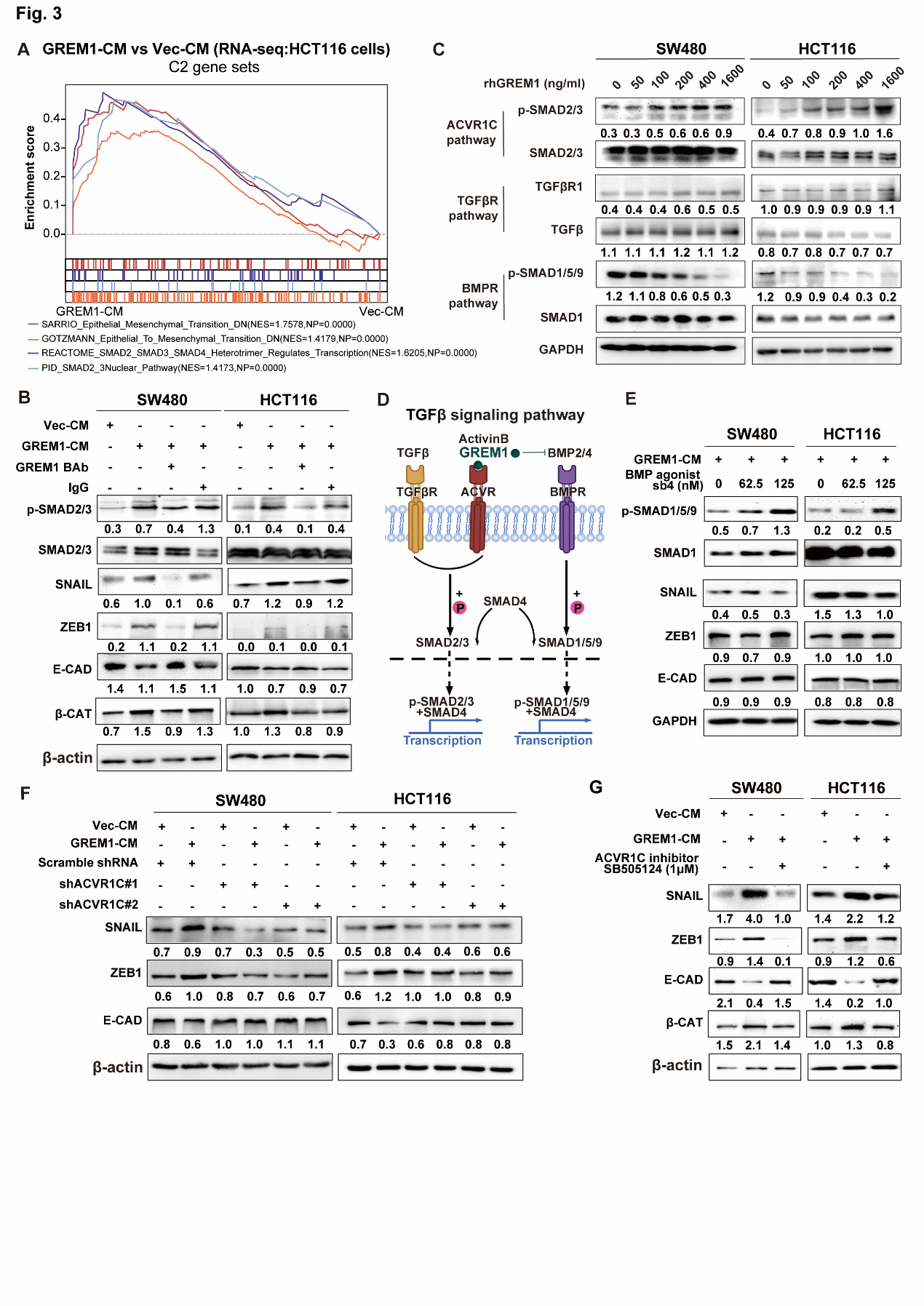

**Figure 4**

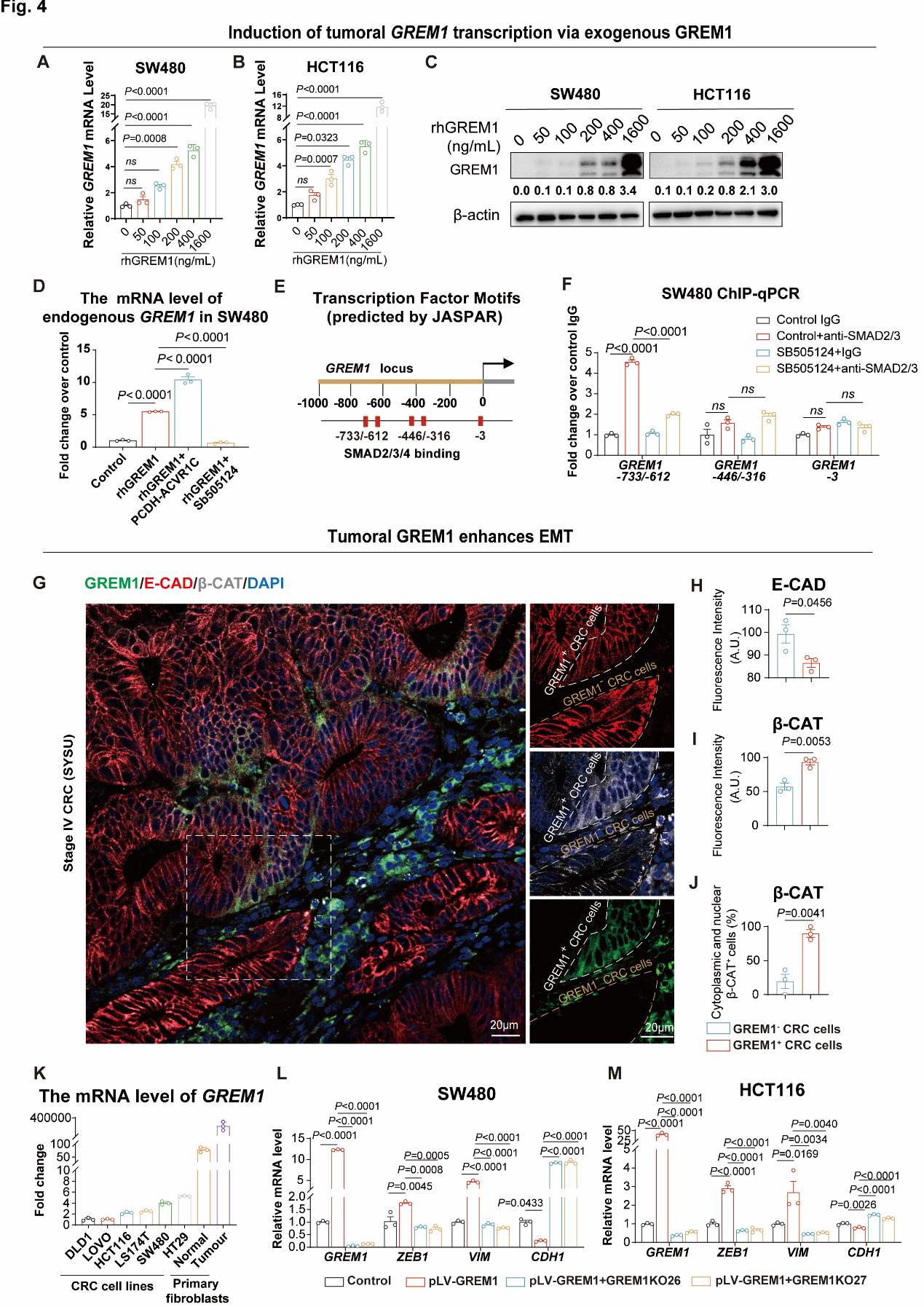

**Figure 5**

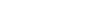

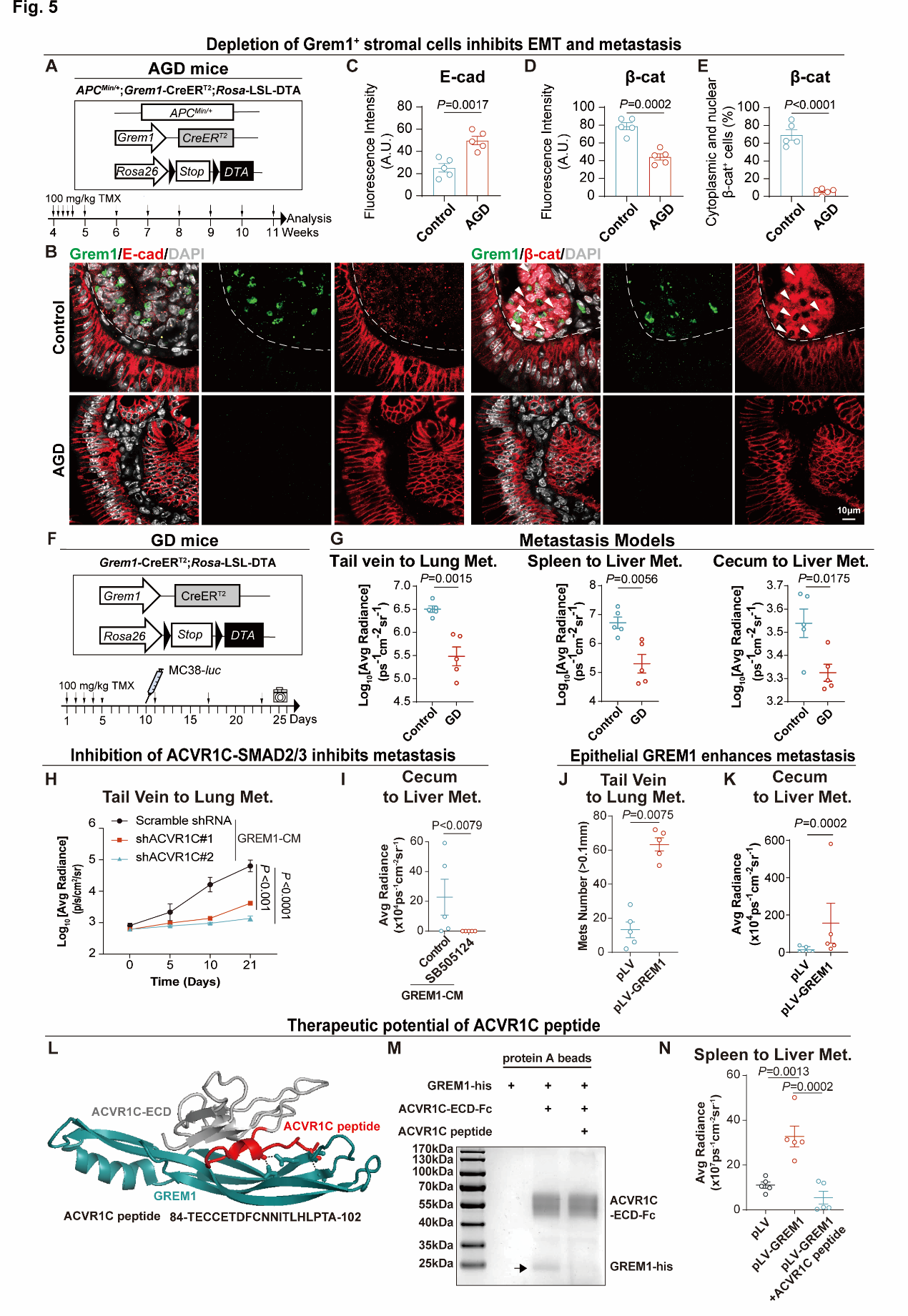

**Figure S1**

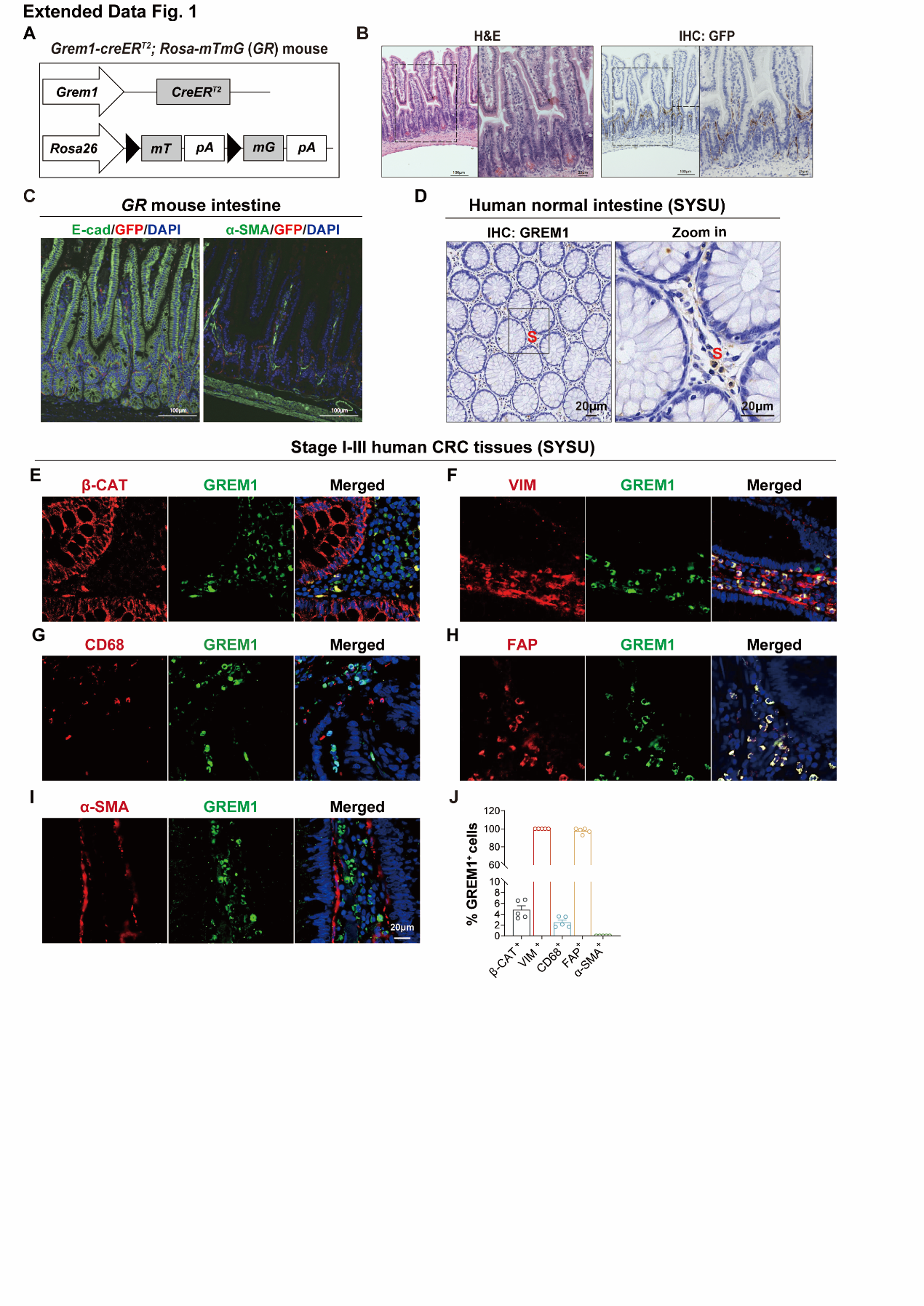

**Figure S2**

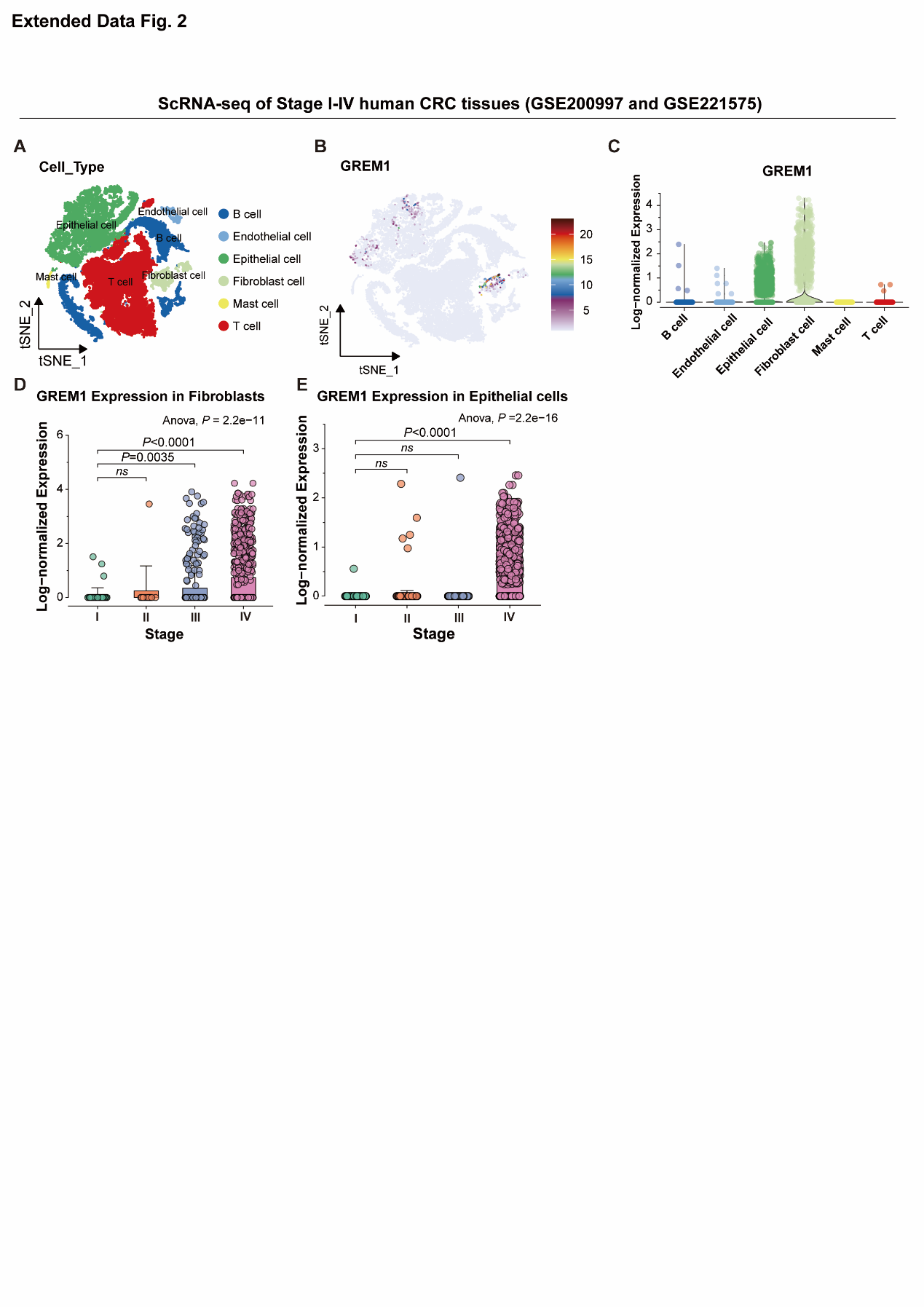

**Figure S3**

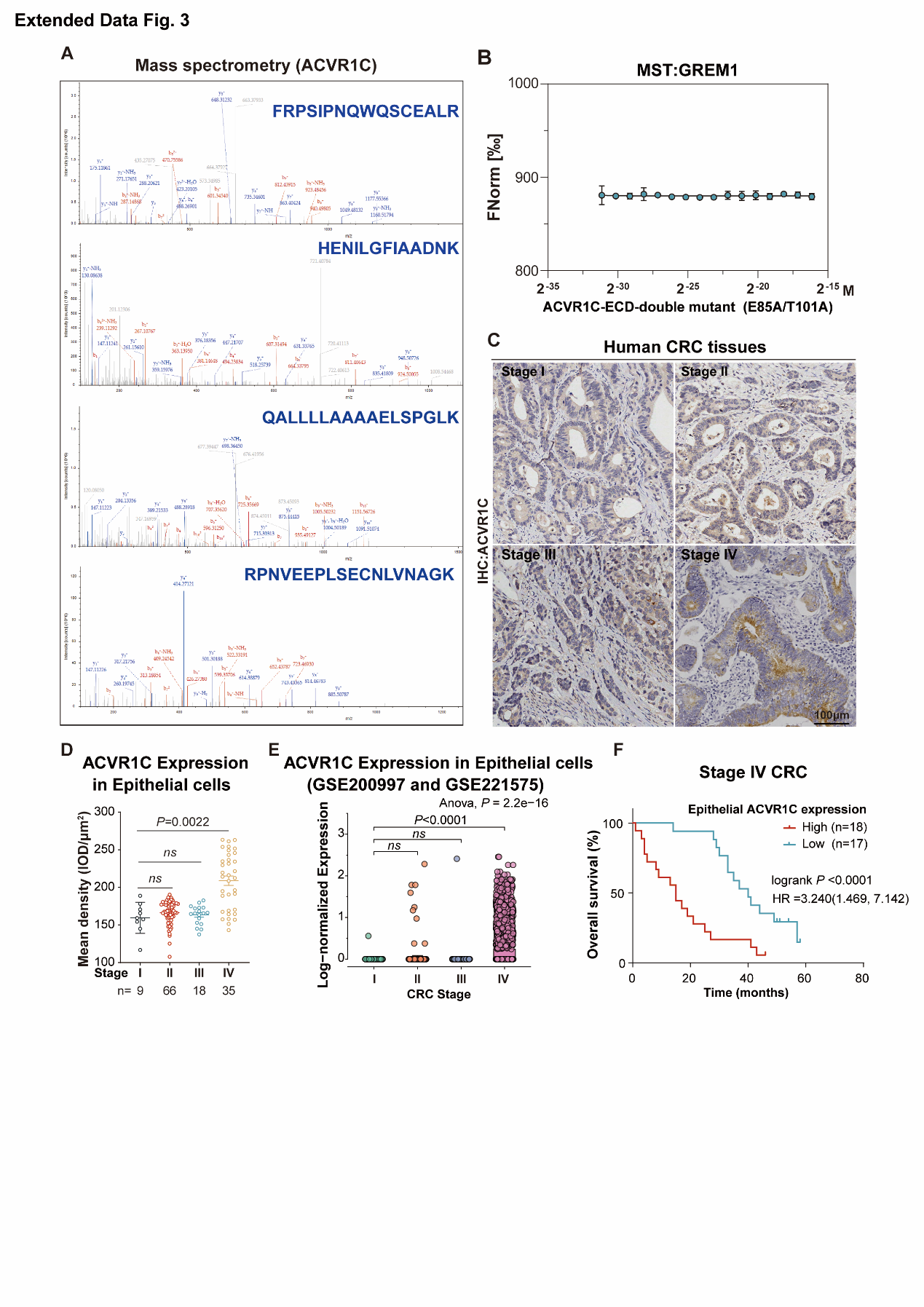

**Figure S4**

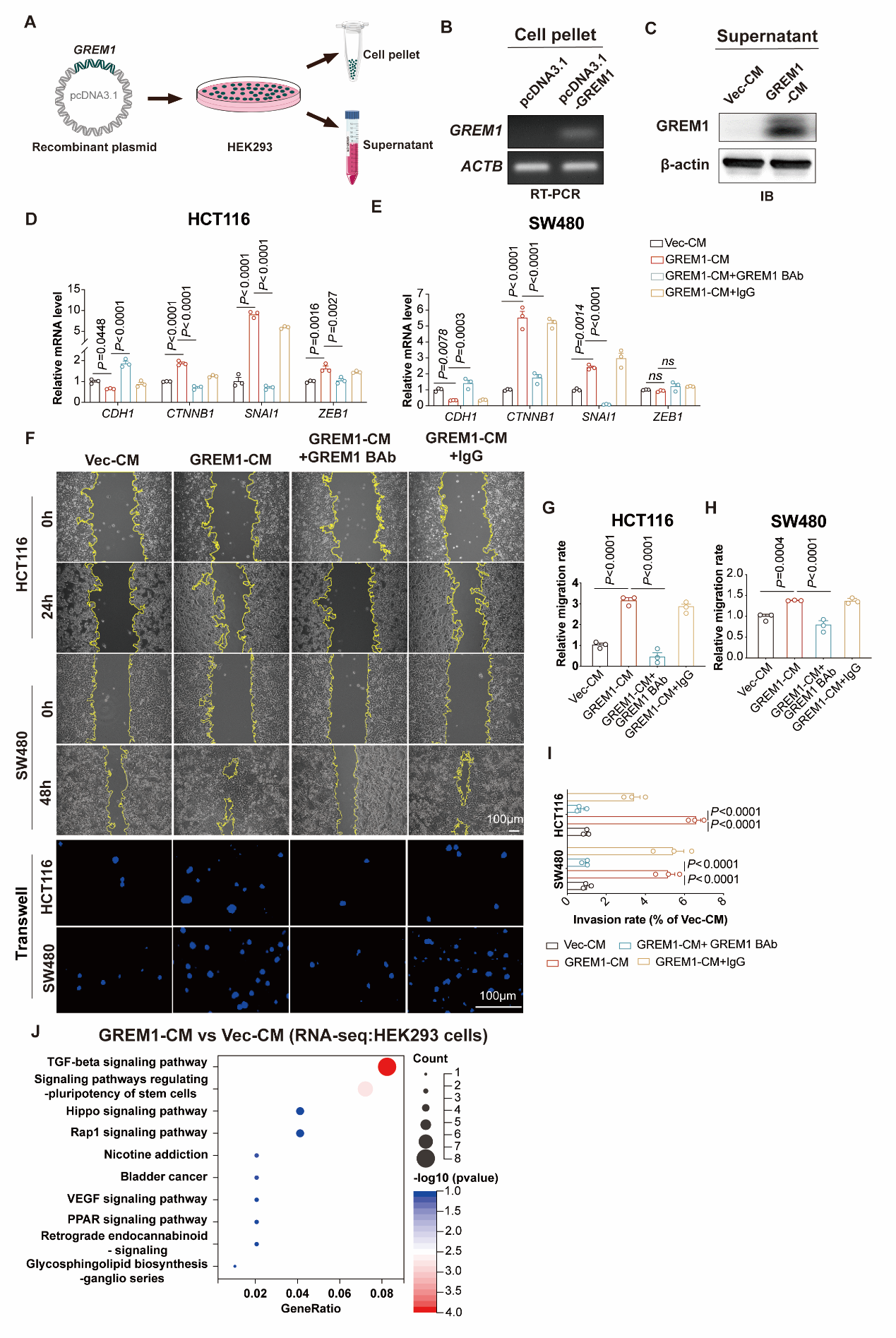

**Figure S5**

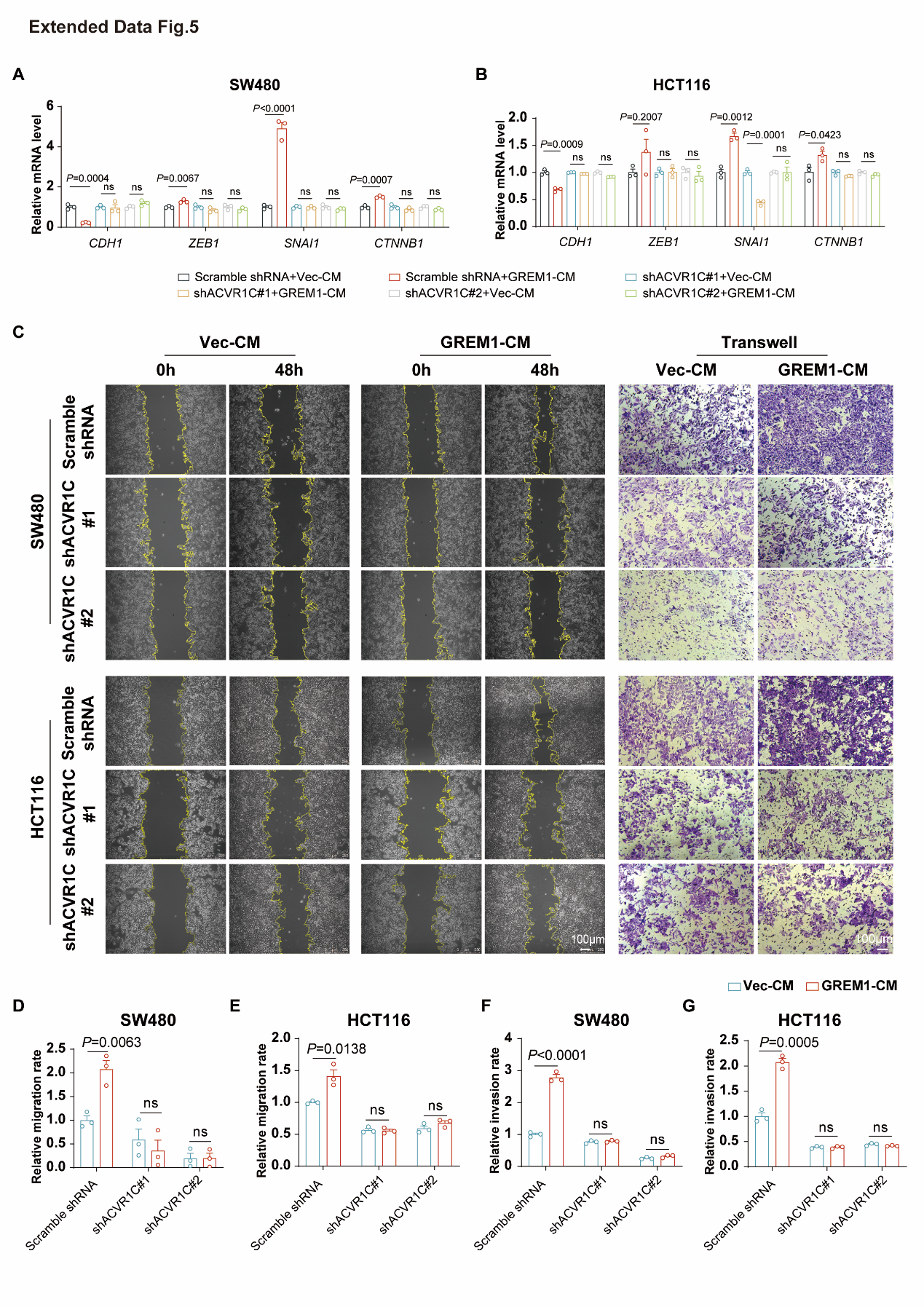

**Figure S6**

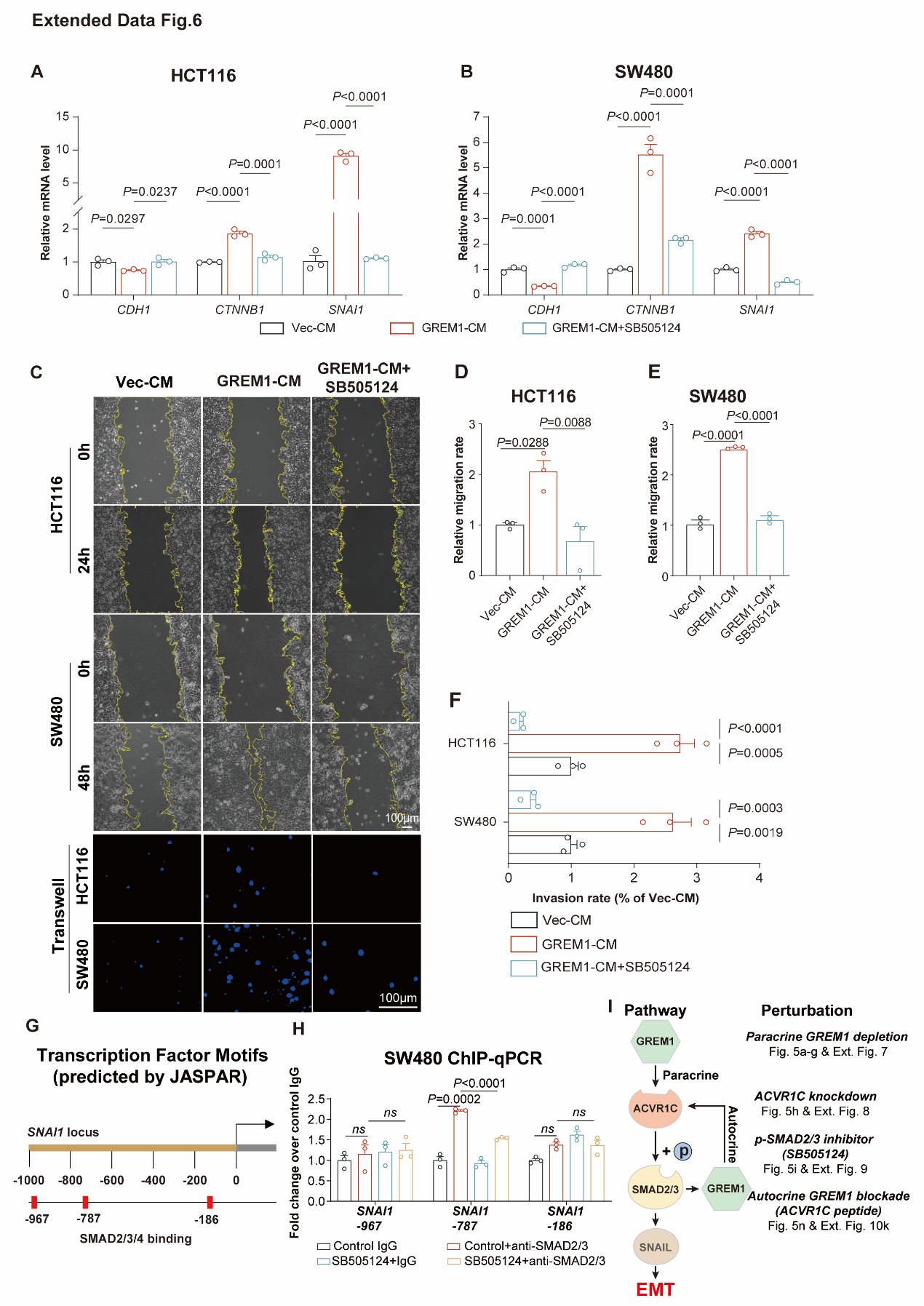

**Figure S7**

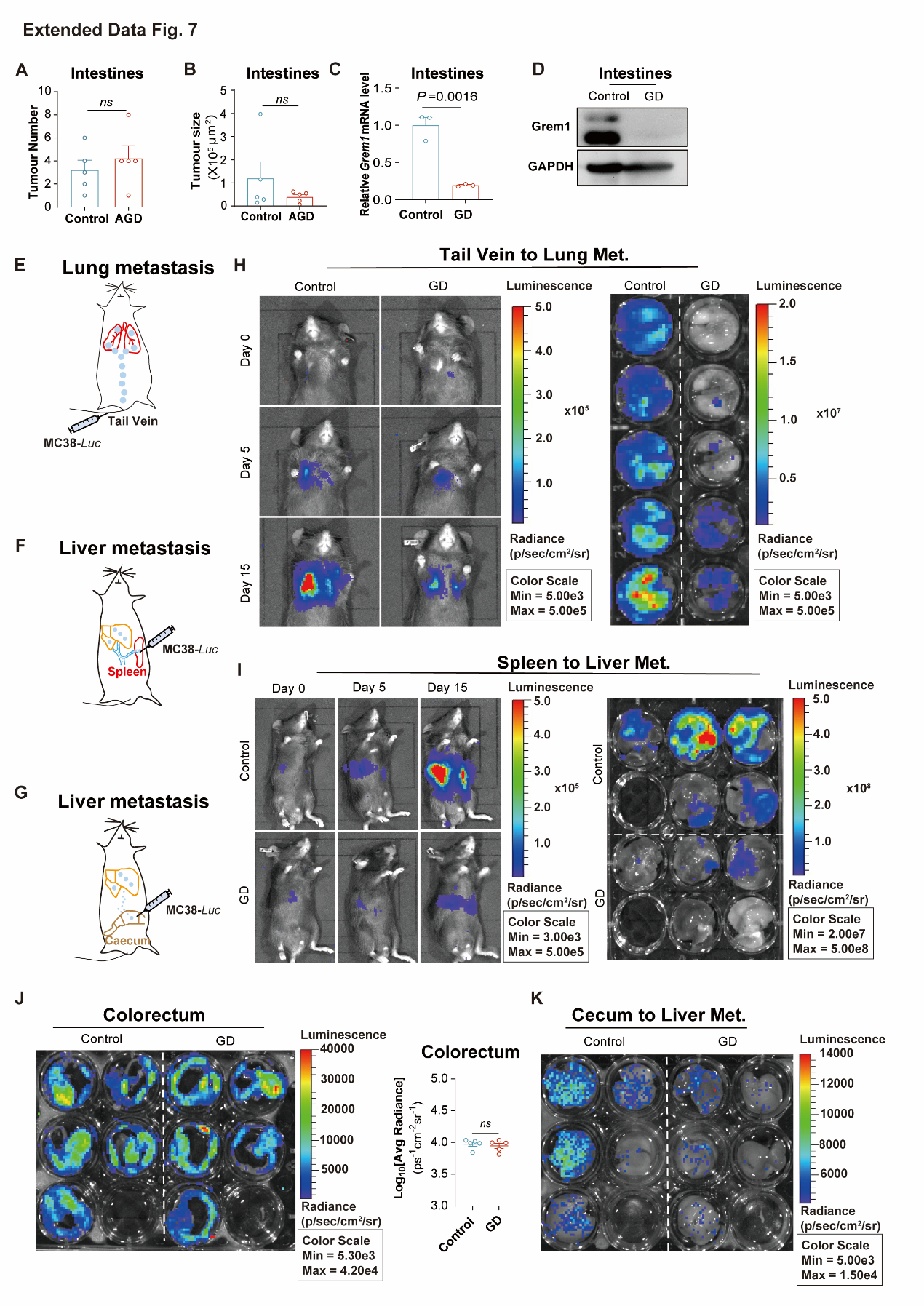

**Figure S8**

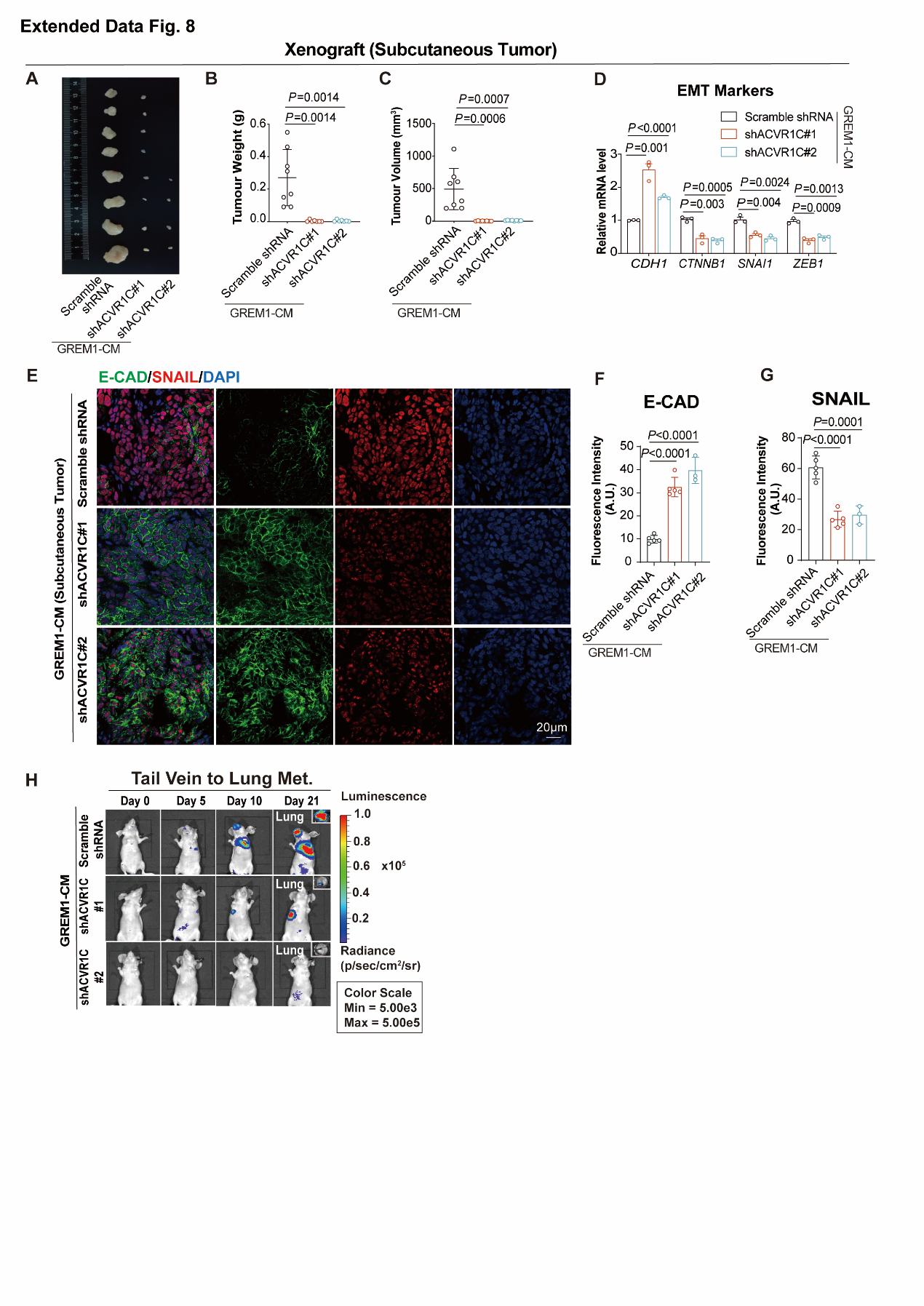

**Figure S9**

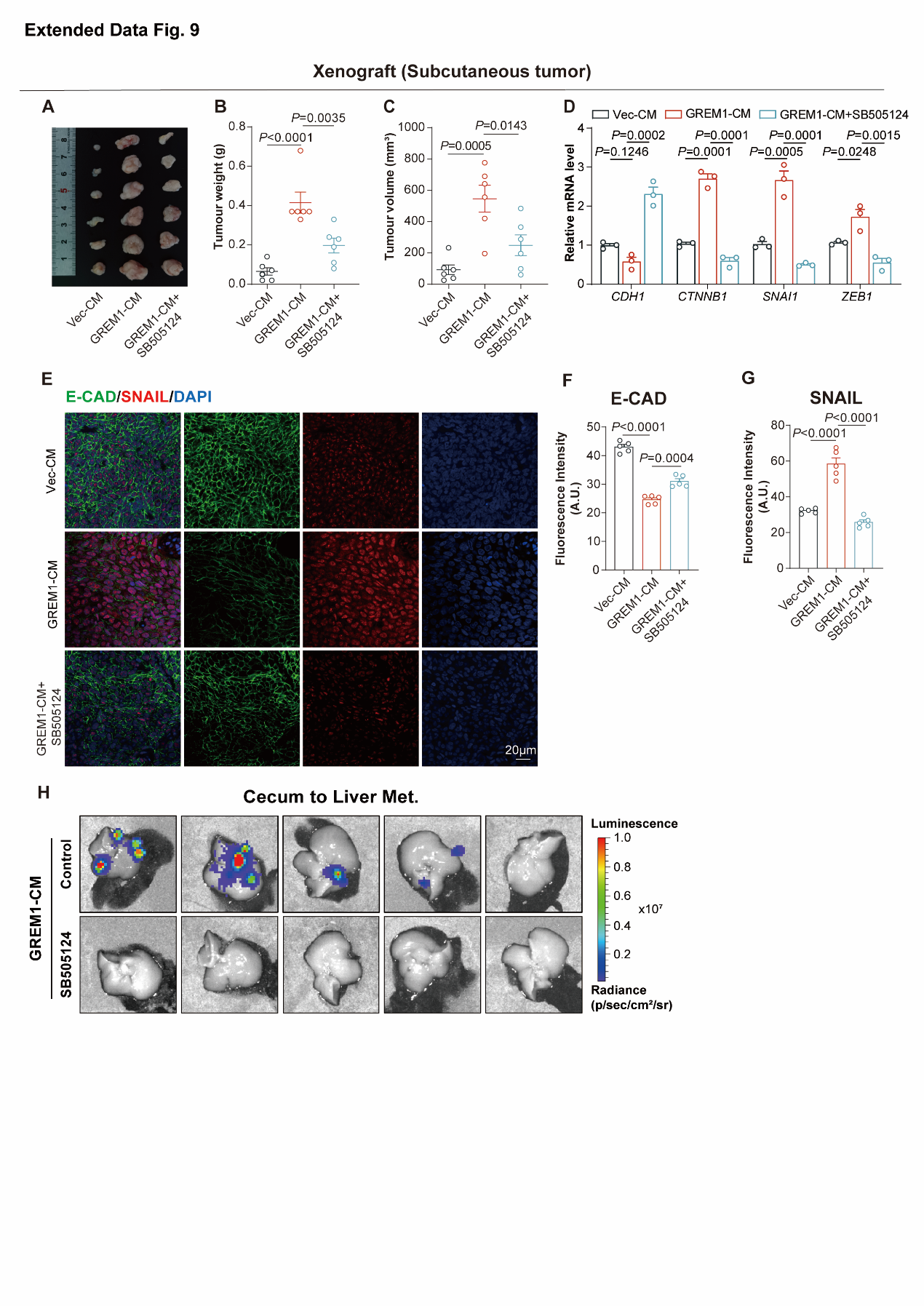

**Figure S10**

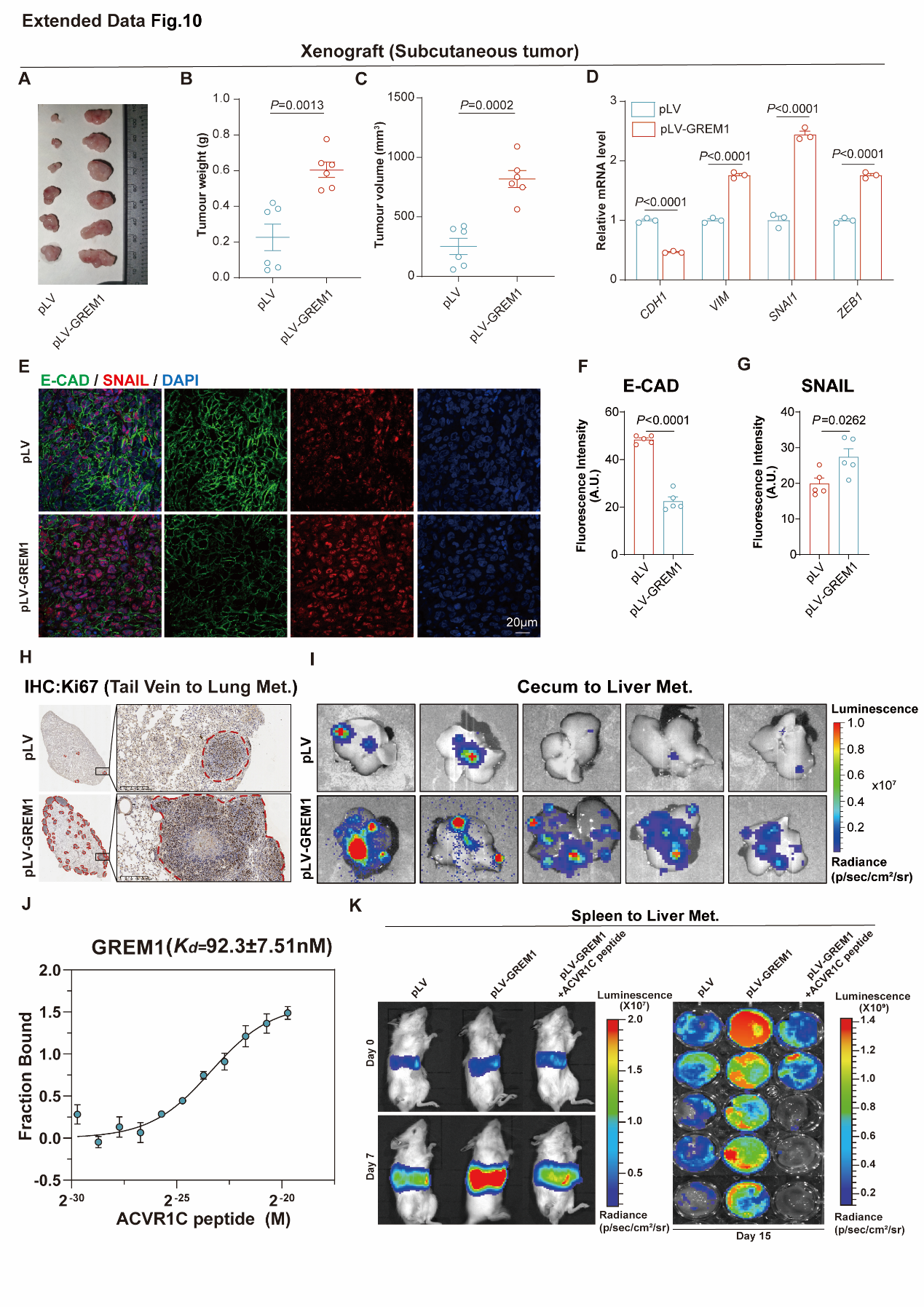

**Figure S11**

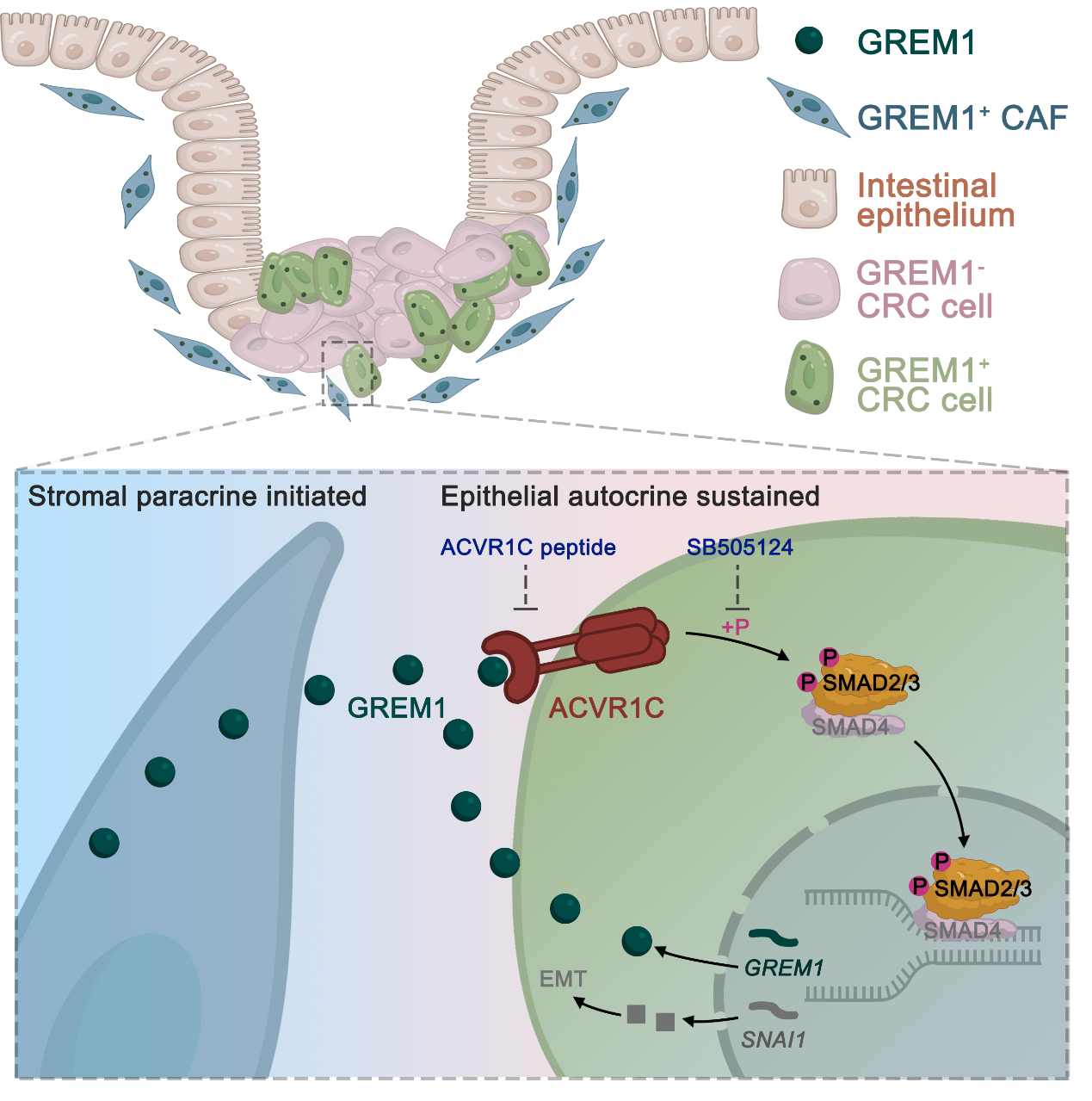

**SUPPLEMENTAL TABLES**

**Table S1. The sequences of RT–qPCR primers used in the study.**

| **Target** | **Species** | **Forward primer sequence (5'–3')** | **Reverse primer sequence (5'–3')** |
| --- | --- | --- | --- |
| *Grem1* | mouse | AGGTGCTTGAGTCCAGCCAAGA | TCCTCGTGGATGGTCTGCTTCA |
| *GREM1* | human | TCATCAACCGCTTCTGTTACGGC | CAGAAGGAGCAGGACTGAAAGG |
| *CDH1* | human | GAGTGCCAACTGGACCATTCAGTA | CACAGTCACACACGCTGACCTCTA |
| *CTNNNB1* | human | GTGACGTTGACATCCGTAAAGA | GCCGGACTCATCGTACTCC |
| *SNAI1* | human | GCTCCCTCTTCCTCTCCATACC | GGCAAGTTGATTGGAGGGATG |
| *ZEB1* | human | TTACACCTTTGCATACAGAACCC | TTTACGATTACACCCAGACTGC |
| *VIM* | human | AAGACGGTTGAAACTAGAGATGGAC | TGCTGGTAATATATTGCTGCACTGA |
| *ACVR1C* | human | ACTTGTGCCATAGCGGACTTA | GGTTCCCACTTTAGGATTCTGAG |
| *GAPDH* | human | AGAAGGCTGGGGCTCATTTG | AGGGGCCATCCACAGTCTTC |

**Table S2. The sequences of ChIP–qPCR primers used in the study.**

| **Target** | **Forward primer sequence (5'–3')** | **Reverse primer sequence (5'–3')** |
| --- | --- | --- |
| *GREM1*–*733/*–*612* | TCGCCCAGTGGCTGTTTT | GTTAACCTGGCGGGCTCG |
| *GREM1*–*446/*–*316* | CAGAGGGGAAGAATGGCC | GAGGGAAGAGCGGGAGGA |
| *GREM1*–*3* | GGGGCGGATAGCGGGTCT | AGAGTGACGCGGCGGCCGTGCA |
| *SNAI1*–*967* | CTCCTACGAGGCCCTGGG | CCCGAGGGAAGAAGTGGC |
| *SNAI1*–*787* | GGAAGCTGCTCTCTAGGAGTTACTC | GCAAAGGGAAGTGTGCTTTGGTG |
| *SNAI1*–*186* | CCGGACAGCCCCAGCAC | CGGAGGCTCGTCTCCGC |
